# In-cell destabilization of a homo-dimeric protein complex detected by DEER spectroscopy

**DOI:** 10.1101/2020.03.27.011510

**Authors:** Yin Yang, Shen-Na Chen, Feng Yang, Xia-Yan Li, Akiva Feintuch, Xun-Cheng Su, Daniella Goldfarb

**Affiliations:** Department of Chemical and Biological Physics, Weizmann Institute of Science, Rehovot 76100, Israel; State Key Laboratory of Elemento-organic Chemistry, College of Chemistry, Collaborative Innovation Center of Chemical Science and Engineering (Tianjin), Nankai University, Tianjin 300071, China

## Abstract

The complexity of the cellular medium can affect proteins’ properties and therefore in-cell characterization of proteins is essential. We explored the stability and conformation of BIR1, the first baculoviral IAP repeat domain of X-chromosome-linked inhibitor of apoptosis (XIAP), as a model for a homo-dimer protein in human HeLa cells. We employed double electron-electron resonance (DEER) spectroscopy and labeling with redox stable and rigid Gd^3+^ spin labels at three protein residues, C12 (flexible region), E22C and N28C (part of helical residues 26–31) in the N-terminal region. In contrast to predictions by excluded volume crowding theory, the dimer-monomer dissociation constant *K*_D_ was markedly higher in cells than in solution and dilute cell lysate. As expected, this increase was recapitulated under conditions of high salt concentrations given that a conserved salt bridge at the dimer interface is critically required for association. Unexpectedly, however, also the addition of a crowding agent such as Ficoll destabilized the dimer, suggesting that Ficoll forms specific interactions with the monomeric protein. Changes in DEER distance distributions were observed for the E22C site, which displayed reduced conformational freedom in cells. Although overall DEER behaviors at E22C and N28C were compatible with a predicted compaction of disordered protein regions by excluded volume effects, we were unable to reproduce E22C properties in artificially crowded solutions. These results highlight the importance of in-cell DEER measurements to appreciate the complexities of cellular *in vivo* effects on protein structures and functions.

## Introduction

It is well known that intracellular environments affect the structural properties of proteins via macromolecular crowding, spatial confinements and different types of cellular interactions. In turn, the stabilities, conformational equilibria and rates of reactions of proteins in the cell may differ from isolated proteins’ behavior in test tubes (1–7). In this context, a fifth level of protein structural organization called quinary structure was introduced (1, 4, 6). Quinary structure denotes the combined effects of cellular environment on individual proteins and it includes the sum of all interactions that a protein experiences in cells. These appear to have co-evolved with protein surface properties to ensure optimal functionality (8). Extensive theoretical (6, 9–11) and experimental efforts have been directed towards understanding and elucidating quinary structure effects for different proteins (see recent reviews(1, 4, 6, 12, 13). Cellular macromolecular crowding described as hard-sphere excluded-volume effects that are entropic in nature favor compact protein conformations, thus acting as stabilizing folded proteins and enhancing protein-protein association (14). Enthalpic contributions via weak attractive or repulsive interactions play equally important roles and can add to or counteract excluded volume effects (9, 15–20). To understand the combined effects of the cellular milieu on protein structure, dynamics, stability, and interactions, in-cell measurements are often combined with *in vitro* measurements under controlled conditions of artificial crowding. Together, these approaches serve to rationalize observed differences between protein behaviors in test tubes and in cells. The number of genuine in-cell studies is small because of the limitations the relevant biophysical methods capable of providing such information face when applied to whole cells. Accordingly, cell lysates are often used as surrogate models but they often fail to recapitulate in-cell effects (6, 21).

Currently most in-cell studies of proteins‘ structure employ nuclear magnetic resonance (NMR), or fluorescence spectroscopy(5, 22), each method having its advantages and limitations. In-cell FRET (Förster energy transfer) is effective for monitoring protein-protein interactions, determining dissociation constants and identifying conformational changes of appropriately fluorophore-labeled target proteins (23–25). In addition, fluorescence correlation spectroscopy (FCS) can be used determine dissociation constants of protein oligomers in cells (26). These techniques however, do not provide structural information on the individual protein-residue level. Recent progress in in-cell NMR spectroscopy has shown that commonly used structural restraints, including nuclear Overhauser effects (NOE) and residual dipolar couplings (RDC) (27–29) as well as paramagnetic restraints such as pseudo-contact shifts (PCS) (30, 31), can be exploited to determine entire protein structures in intact cells. Nonetheless, many proteins display poor in-cell NMR signal qualities (line broadening) due to restricted motions and transient interactions with cellular components (3, 32, 33). In-cell studies using NMR and fluorescence techniques reported on protein folding and stability in cells (17, 24, 34–40), structural changes of disordered proteins (41–43) and, to some extent, on protein association e and the stabilization of oligomeric protein forms (23, 25, 26).

EPR (electron-paramagnetic resonance)-based double electron–electron resonance (DEER, also called PELDOR) spectroscopy has been suggested as an attractive alternative method to interrogate protein structures in cells (21, 42, 44–55). DEER provides distance distributions between two spin labels attached to well-defined positions in proteins (56) and can thus be used to probe proteins conformations and to deduce information about oligomeric protein states (57, 58). DEER measurements are typically carried out on frozen solutions due to relaxation-time limitations and, therefore, are limited in the dynamic information they can provide. Nonetheless, the widths of distance distributions report on degrees of conformational freedom at the time of freezing. To date, most of the published in-cell DEER studies have focused on establishing the methodology, on optimizing spin-label performance and on demonstrating that such measurements are indeed feasible (46, 47, 49–53, 55). By contrast, here we address a fundamental biological question by interrogating the stability of a homo-dimeric protein complex in human HeLa cells and by comparing our findings to results that we obtain in dilute cell lysates and under isolated buffer conditions.

The protein studied is the first baculoviral IAP repeat (BIR1) domain of X-chromosome-linked inhibitor of apoptosis (XIAP). XIAP is a multidomain protein that is directly involved in caspase inhibition and is therefore a potential target for cancer therapy (59, 60). XIAP also participates in receptor signaling, cell division, ubiquitin ligation, and cellular copper homeostasis (61, 62). It harbors three zinc-binding BIR domains (BIR1–BIR3), the second and third of which are responsible for interactions with caspase (63–66). The structures of the three BIR domains have been determined by NMR spectroscopy and X-ray crystallography (63–69) and BIR1 has been found to exist as a stable homodimer *in vitro* (63, 65, 66). Furthermore, interaction of BIR1 dimer with the TAK1-binding protein TAB1 was reported to be essential for activation of the NF-κB signaling pathway (69). Residues 1–19 of N-terminal BIR1 dimers are not observed in the X-ray structure (68, 69) (Fig. 1a), and they are highly flexible as determined by NMR (67). The main interactions between BIR1 monomers are mediated by a salt bridge between D71 in one monomer and R72 in the other; breaking this salt bridge produces the monomeric protein in solution without major structural changes in domain architecture (67). In a previous study, we reported that in-cell NMR spectra of ^15^N-labeled BIR1 either overexpressed in *E. coli*, injected into *Xenopus laevis* oocytes, or transduced into HKT293 cells showed no observable NMR signals (67), suggesting that cellular interactions broadened BIR1 resonances beyond detection.

**Figure 1.**
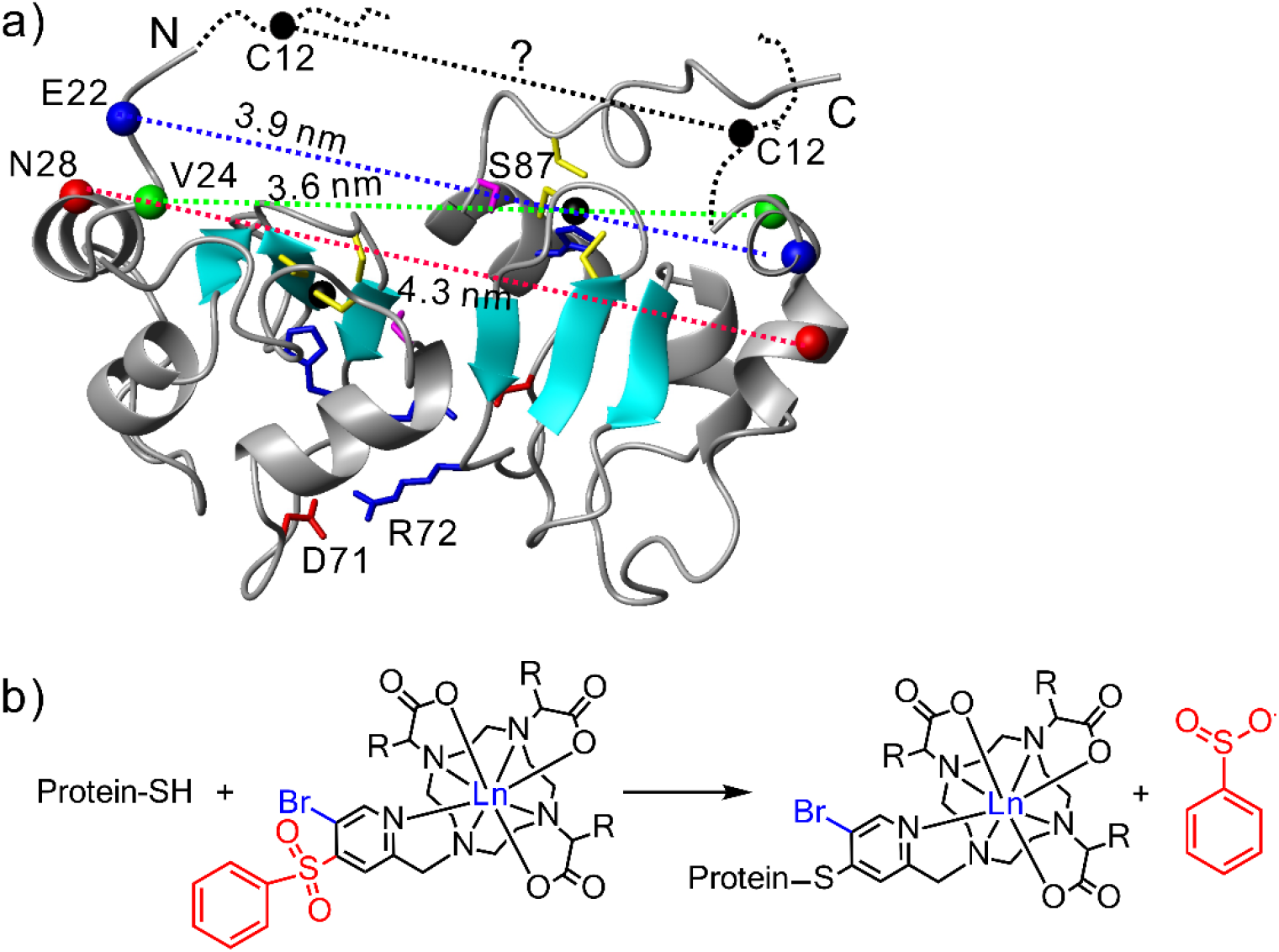
(a) Structural representation of dimeric BIR1 (PDB code: 2QRA(68)). N-terminal residues (1–19) are indicated by black dashed lines as they are not observed in the crystal structure. In the dimer, distances between Cα atom pairs of E22, V24, and N28, mutated to Cys for spin labeling, are indicated by colored dashed lines. Side chains of S87, C63, C66, C90, and H83 of the zinc finger motif are shown as sticks. Zinc ions are indicated as black spheres. The main interaction at the dimer interface is a salt bridge between D71 and R72. These residues are shown as sticks and labeled. (b) Chemical reaction of site-specific BIR1 labeling with BrPSPy-DO3MA-Ln (R = CH_3_) or BrPSPy-DO3A-Ln (R = H) for DEER and NMR measurements.

In this work, and using BIR1 as a homodimeric model complex, we specifically asked: (i) How does the cellular environment affect the stability of the BIR1 dimer in comparison to dilute cell lysate and isolated in a buffer solution? (ii) Can we mimic in-cell behaviors under appropriate buffer conditions? (iii) Does the BIR1 domain undergo structural changes in cells? We addressed these questions by W-band (95 GHz) DEER measurements of site-specifically engineered BIR1 that we labeled with rigid paramagnetic Gd^3+^ tags (52, 53) via redox stable thioether bond (C-S) formation (Figure 1b). Our results show that the monomer-dimer dissociation constant, *K*_D_, is higher in cells than in dilute lysate and buffer. This indicates that the intracellular environment of HeLa cells destabilizes the BIR1 dimer. We were able to reproduce this effect under high salt conditions *in vitro* and in artificially crowded solutions. We additionally found that segmental motions of N-terminal BIR1 residues experienced greater attenuations in intact cells than in cell lysate and buffer. We were unable to reproduce these changes in the presence of salt or crowding. We discuss these results in light of the expected behavior by classic excluded volume theory and with respect to earlier experimental observations pointing towards destabilizing quinary structure effects.

## Results

### Selection of spin labeling sites

In this workwe used two spin labels, BrPSPy-DO3A-Gd^3+^ and BrPSPy-DO3MA-Gd^3+^ (52, 53) (see Fig. 1b), referred to as **GdI** and **GdII** hereafter. The linkage of these spin labels to the protein is rigid, which minimizes contributions of inherent spin label dynamics to distance distributions, thereby increasing the precision of measured distance information. While both spin labels produce similar distance distributions, **GdI** yields better sensitivity than **GdII** because of its narrower EPR spectrum (52). To probe N-terminal part of BIR1, we spin-labeled residues C12, E22C and N28C, which are far from the dimer interface in order to minimize destabilizing the dimer or perturbing its three-dimensional (3D) structure. These sites are expected to exhibit increasing flexibilities according to C12>E22>N28 based on the known BIR1 structure (see Fig. 1a). C12 resides in the unstructured part of the N-terminal region, whereas residue E22 is located near the end of the flexible N-terminal segment. N28 is located in the middle of the first BIR1 α-helix (68). We ligated **GdI** and **GdII** to C12 in wild type (WT) BIR1 and to the cysteins in the C12A/E22C and C12A/N28C generated mutants. In these constructs, we mutated native C12 to alanine, such that each monomer contained only one labeling site (see Methods section in SI for details). In addition to C12, BIR1 harbors three additional cysteine residues, C63, C66, and C90, that constitute a zinc binding site, which are critical for the stability of the BIR1 domain fold. Zinc removal in the presence of excess EDTA results in unfolding (67). Mass spectrometry confirmed that each protein construct was labeled at the expected, single site (Fig. S1, SI) and we found that both BrPSPy-DO3MA-Ln and BrPSPy-DO3A-Ln tags do not compete with the zinc binding site.

Using NMR, we confirmed that ligation of WT BIR1 or its cysteine mutants with BrPSPy-DO3MA-Ln (Ln = Gd^3+^, Dy^3+^, Tm^3+^, or Y^3+^) or **GdI** did not change the overall architecture of the domain (Fig. S2–S3, SI) and that zinc was properly bound to C63, C66, and C90 (Fig. S1, SI). We experimentally confirmed the expected decrease in protein motions along the C12, E22C and N28C series by paramagnetic NMR measurements (Fig. S3, SI), wherePCS increased as follows :C12<< E22C< N28C (Fig. S4, SI). Because each BIR1 monomer contained one spin label, only dimers should contribute DEER modulations. To verify this we carried out W-band DEER measurements on monomeric D71N/R72E C12-**GdII** and on D71N/R72E/C12A/N28C- **GdII** and as expected, the DEER traces did not reveal modulations in solution (Fig. S5, SI). Next, we analyzed WT BIR1, C12A/E22C and C12A/N28C, labeled with **GdI** and **GdII** in frozen solutions, in HeLa cell lysates and intact Hela cells. For the latter, we delivered spin-labeled BIR1 into the cell by electroporation(42, 70). Homogenous intracellular distribution of BIR1 was confirmed by microscopy imaging of C12-ATTO488 fluorophore-labeled WT BIR1 (Fig. 2a). Examples of echo-detected EPR (ED-EPR) spectra of C12A/E22C–**GdI** in buffer, cell lysates and cells are shown in Fig. 2a. ED-EPR spectra of all the other samples labeled with **GdI** are depicted in Fig. S6, SI. All spectra in in cells and cell lysates displayed signals of endogenous Mn^2+^ (53, 54). Corresponding echo decay measurements are shown in Fig. S7, SI.

**Figure 2.**
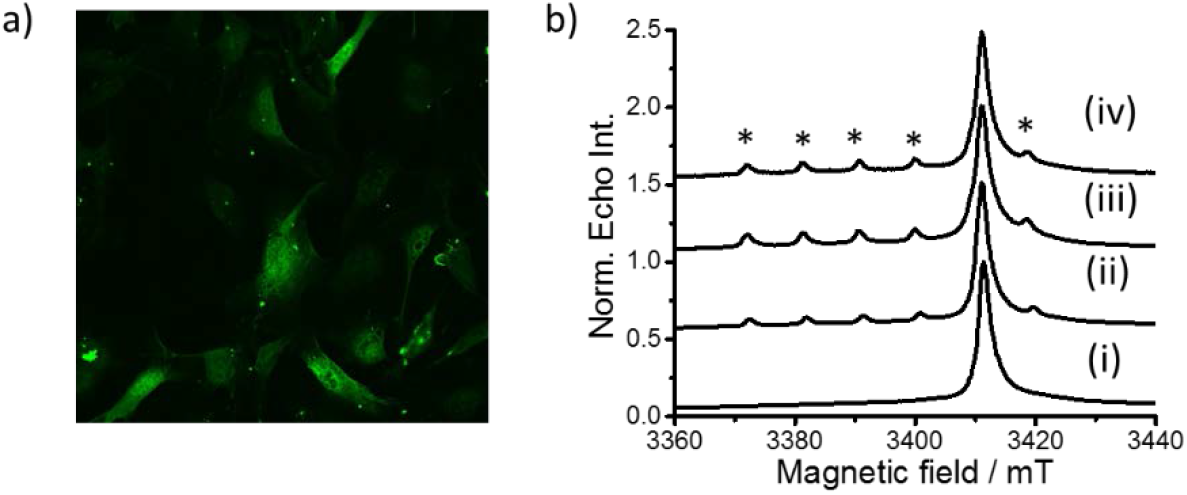
(a) Fluorescence microscopy image of WT C12–ATTO488 BIR1 after delivery into HeLa cells by electroporation. (b) Central transition regions of W-band ED-EPR spectra of C12A/E22C–**GdI** BIR1 in (i) buffer (200 μM BIR1), (ii) HeLa cell lysate (30 μM BIR1), (iii) intact HeLa cells, and (iv) lysed HeLa cells after BIR1 delivery and incubation for 5 h at 37 °C. All spectra were normalized and are shifted along the y axis to ease comparison. Signals marked with asterisks correspond to cellular Mn^2+^.

### Cellular environments affect the conformations of BIR1 N-terminal flexible segments

DEER data from frozen solutions (Fig. 3) confirmed BIR1 dimerization and revealed widths of distance distributions with the trend C12>E22C>N28C. These results are consistent with the degrees of ligation site dynamics determined by NMR spectroscopy and with expectation from the BIR1 crystal structure (Fig. 1a). Distance distributions in HeLa cell lysates (Fig. 3) were s to distributions measured in buffer with close modulation depths (λ as defined in Fig. 3a). These results suggested that dimer formation was favored in cell lysates despite significantly lower BIR1 concentrations (30 μM versus 200 μM, respectively). Distance distributions obtained for E22C and N28C mutants agreed reasonably well with those predicted with MtsslWizard from the BIR1 crystal structure (71) (Fig. 3).

**Fig. 3.**
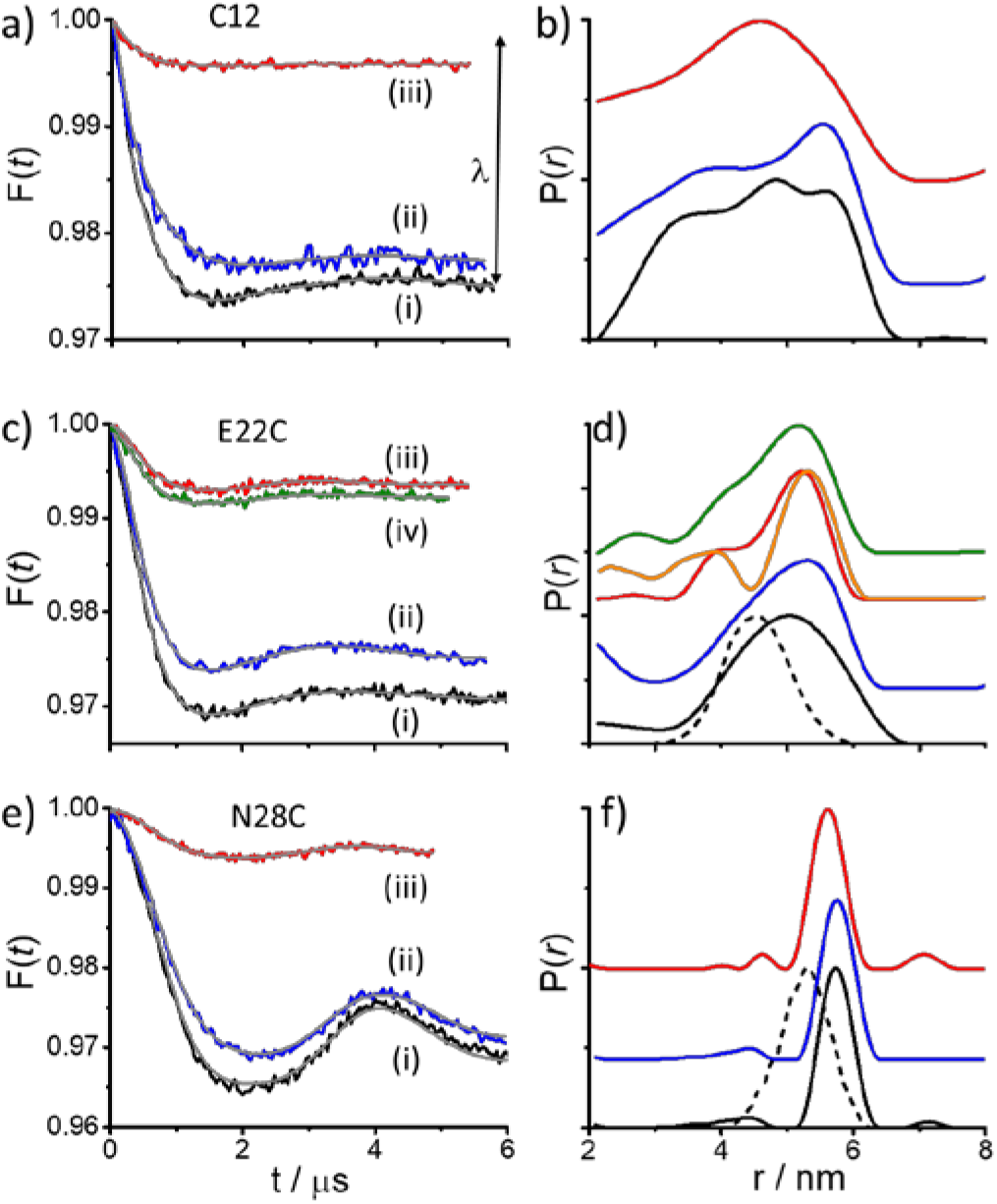
(a, c, e) W-band DEER data after background corrections (form factors, F(t)), fitting (gray lines)), and respective distance distributions (b, d, f). Results for WT C12–**GdI** (a, b), C12A/E22C–**GdI** (c, d), and C12A/N28C–**GdI** (e, f) in buffer (i, 200 μM BIR1, black), primary lysates (ii, 30 μM BIR1, blue), intact (iii, red) and lysed cells (iv, green). Raw DEER data are in SI Fig. S8–S10. Distance distributions for C12A/E22C–**GdII** are shown in orange (d) (see also Fig. S9, SI). Distance distributions predicted by MtsslWizard (71) (PDB code: 2QRA (40)) are indicated by dashed lines (d, f). The arrow in (a) depicts the modulation depth λ for the solution sample. Shown distance distributions are normalized. We note that by breaking the cell membrane, the effective biomacromolecular concentration reduced to 2-3 times lower than the intact cells.

We measured broad distance distributions for WT BIR1 C12–**GdI** in HeLa cells, which indicated that this part of the N-terminal segment remained flexible and unstructured. We also detected a narrow in-cell distance distribution for C12A/N28C–**GdI**, with small shifts of respective maxima (−0.2 nm) for the in-cell sample (Fig. 3e,f). Similar results were obtained for **GdII**-labeled BIR1 (Figs. S8–S10, SI). By contrast, C12A/E22C-**GdI** BIR1 revealed narrower distance distributions in cells than in lysates and buffer for both versions of spin-labeled BIR1 (i.e. **GdI** and **GdII**) (Fig. 3d). We confirmed the reproducibility of this observation by additional triplicate measurements of **GdII** BIR1 constructs (Fig. S9, SI). To further rationalize this discrepancy, we performed DEER measurements on C12A/E22C-**GdI**transduced HeLa cells that we lysed after 5 h of incubation at 37 °C in the EPR sample tube (see Fig. 3c,d). In turn, we measured a broad distance distribution similar to primary lysates and in buffer. Surprisingly, however, the modulation depth did not change and remained as low as in intact cells. The main difference between these lysed in-cell samples and primary lysate specimens relates to the relative amounts of BIR1 versus total cellular components, which are 5-7 time lower for the latter (see SI experimental section). Therefore, our results demonstrated that intact cellular environments and genuine intracellular crowding conditions affected the structural properties of the BIR1 domain around the region spanning E22C. Next, we set out to recapitulate these effects with artificial crowding agents such as Ficoll and bovine serum albumin (BSA). Unexpectedly, we were unable to achieve similar degrees of distance-distribution-narrowing (Fig. S11-12, SI). We also failed to induce such effects upon addition of a two-fold molar excess of trifluoroethanol (TFE), which is known to stabilize alpha-helicity(72). By contrast, addition of a ten-fold excess of TFE abolished the modulations all together (Fig. S13, SI).

To determine the extent by which the HeLa cytoplasm modulates the dynamics of E22C proximal regions in BIR1, we introduced additional single cysteine mutations at K19 and V24 for spin-labeling with **GdI**. We measured rather broad distance distributions for both mutants in buffer, similar what we determined for C12A/E22C-**GdI** (Fig. 3d). Furthermore, the maximum distance distribution for C12A/V24C–**GdI** was in good agreement with crystal structure predictions (Fig. S14-S15, SI). Distance distributions of C12A/K19C–**GdI** and C12A/V24C–**GdI** in-cell samples displayed only subtle differences that we deemed within the experimental error of the method.

### DEER results indicate decreased stability of the BIR1 dimer in cells

Whereas in-cell DEER data of **GdI** spin-labeled BIR1 constructs (i.e. WT, C12A/K19C, C12A/E22C, C12A/V24C, and C12A/N28C) revealed the presence of dimers in intact HeLa cells, all exhibited λ values that were 1/4–1/6 of the corresponding specimens in buffer (Fig. 3, and Fig. S14-S15, SI). We noted similar properties for **GdII** spin-labeled samples (see Fig. S8-S10, SI). These substantial reductions of modulation depths were in stark contrast to results in cell lysates, where λ’s were only marginally reduced. We expected cellular reduction in λ in the cell owing to the presence of Mn^2+^ (53). However, and given the relative differences of Gd^3+^ and Mn^2+^ in-cell signal intensities at the positions of the observe pulses (maximum of the Gd^3+^ signal, see Fig. 2 and Fig. S6, SI), Mn^2+^ effects should be too small to account for the large decrease in λ. Alternatively, we considered that changes in BIR1 monomer-dimer equilibria may cause the observed effect. Low intracellular BIR1 concentrations may indeed shift the equilibrium towards higher amounts of monomeric protein. To address this possibility, we measured dimer dissociation constant (*K*_D_) from DEER modulation depths, known to be proportional to % of dimers in solution (73). DEER experiments on buffer solutions containing different concentrations of C12A/N28C–**GdI** BIR1 (10–400 μM monomer) showed a clear concentration dependence for λ (Fig. 4a, b). Data fitting yielded *K*_D_= 11.3 ± 3.8 μM (Fig. 4c) (details in SI). Importantly, DEER traces produced equal distance distributions in all cases, except for the 10 μM sample, which revealed broader distributions because of poor signal-to-noise ratios (SNR) and the ensuing uncertainties in background decay traces (Fig. S16, SI).

**Fig. 4.**
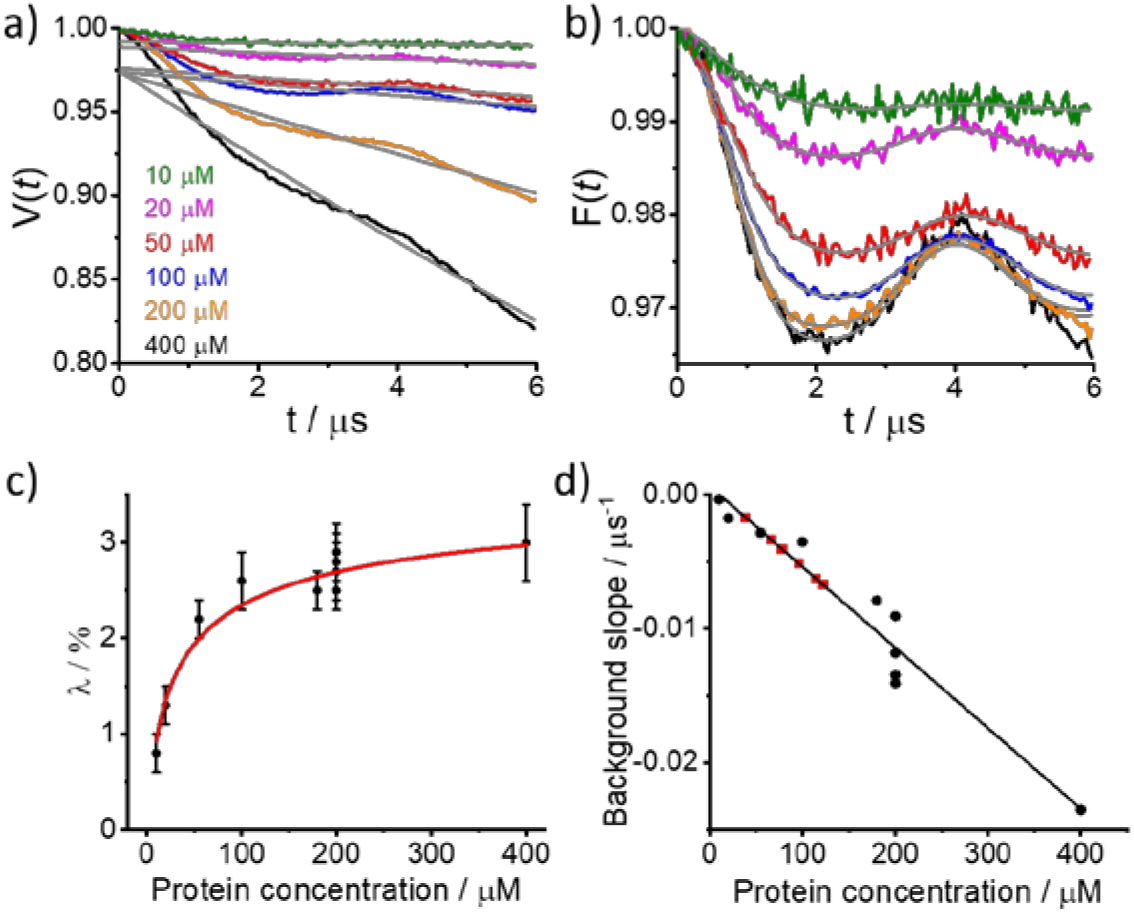
BIR1 dimer dissociation constants from W-band DEER experiments. (a) Primary DEER data on C12A/N28C-**GdI** at indicated concentrations in 20 mM Tris (pH 7.2). Background decay traces shown in gray. (b) DEER form factors after background removal and data fitting (gray lines) obtained with distance distributions shown in Figure S16a, SI. (c) Modulation depths (λ, black symbols) as functions of BIR1 concentrations (monomer) and the fit (red line) to determine *K*_D_. (d) Linear calibration curve with data points (black) from background decay slopes of samples in (a) giving a slope of 6.0 ± 0.53 × 10^−5^ μs^−1^ μM^−1^. Data points of in-cell samples are shown in red. For plots in (c) and (d) solution samples data of all mutants labeled with **GdI** were included.

We also measured the dimer dissociation constant of the C12A/S87A/N28C–**GdI** mutant (Fig. 5). Notably, mutation of S87 to alanine prevents cellular phosphorylation of S87, which, in turn, may impede dimer stability. The motivation for these experiments was the reported phosphorylation of XIAP at S87 by protein kinase C *in vitro* and cells(74) and the lack of a DEER effect in solution for the S87E mutant labeled at C12 with **GdII** (Fig. S5c, SI). The latter suggests that phosphorylation at this site should destabilize the dimer considering that S-to-E mutation is often used to mimic phosphorylation (40). Measured concentration-dependent changes of λ yielded a *K*_D_ of 97.5 ± 22.6 μM, which is considerably larger than for WT BIR1.

**Fig. 5.**
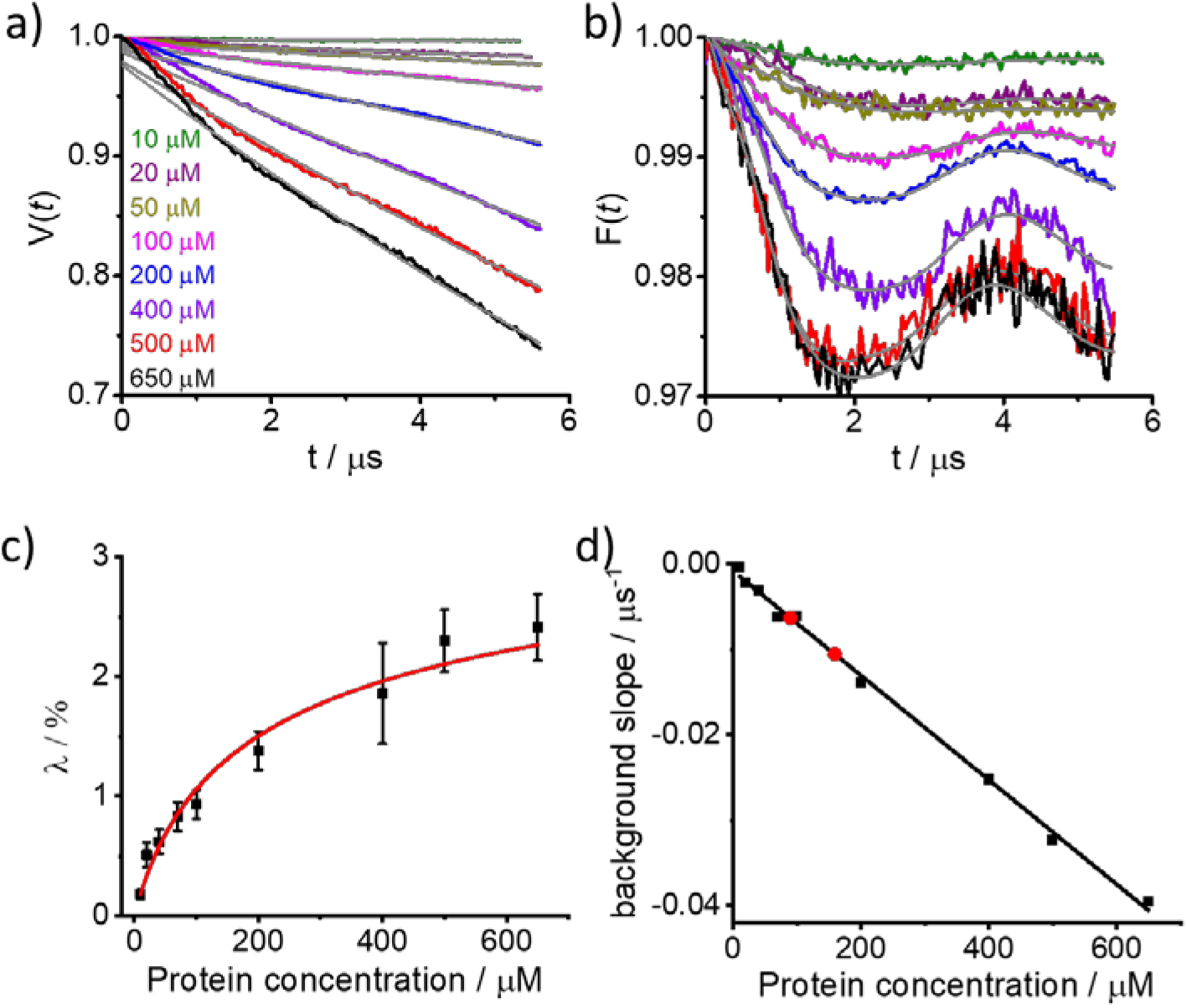
BIR1 S87A/C12A/N28C-**GdI** dissociation constants from W-band DEER measurements. (a) Primary DEER data at indicated protein (monomer) concentrations in 20 mM Tris (pH 7.2) with background decay traces shown in gray. (b) DEER form factors after background subtraction and data fitting (gray lines) with distance distributions shown in Fig. S16b, SI. (c) Modulation depths as a function of BIR1 concentrations and fit (red line) used to the determine *K*_D_. (d) Linear calibration curve with data points (black) from background decay slopes of samples shown in (a), yielding a slope of 6.1 ± 0.13 × 10^−5^ μs^−1^ μM^−1^. Data points of in-cell samples are shown in red.

To substantiate the existence of a BIR1 monomer-dimer equilibrium, we carried out concentration-dependent 2D ^1^H-^15^N hetero-nuclear single quantum coherence (HSQC) NMR experiments on ^15^N isotope-labeled, WT BIR1 (Fig. S17, SI). We obtained a *K*_D_ of 4.7 ± 2.0 μM (at 298 K), which was consistent with size-exclusion chromatography results at different BIR1 concentrations (Fig. S18, SI). The determined *K*_D_ value is smaller than that obtained by DEER, which likely reflected temperature effects between NMR (room temperature) and DEER experiments (freezing point of samples) (75). The presence of spin-labels and 20% glycerol in the latter may have further contributed to the observed differences. Similarly, the S87A BIR1 mutant exhibited a monomer–dimer equilibrium in buffer, as evidenced by size exclusion chromatography (Fig. S18, SI) and NMR spectroscopy (Fig. S19, SI). The *K*_D_ of this mutant, determined by NMR spectroscopy in analogy to WT BIR1, was 20.3 ± 4.6 μM (at 298 K). This value is significantly higher than that of WT BIR1, consistent with our DEER results. This value is also smaller than the one determined by DEER, as discussed earlier. Table I lists measured *K*_D_ values.

**Table 1.** Summary of the *K*_D_ values of BIR1 samples measured by DEER and NMR under various conditions.

| Sample (DEER) | $K_D$ (DEER), $\mu\text{M}$ | Sample (NMR) | $K_D$ (NMR), $\mu\text{M}$ |
| --- | --- | --- | --- |
| C12A/N28C-**Gdl**,<br>frozen solution | $11.3 \pm 3.8$ | WT, solution | $4.7 \pm 2.0$ |
| S87A/C12A/N28C-**Gdl**<br>frozen solution | $97.5 \pm 22.6$ | S87A, solution | $20.3 \pm 4.6$ |
| Frozen cells, all labeled<br>mutants | $126 \pm 40$ | | |

The evaluation of the expected in-cell λ for the *K*_D_ value determined in frozen solutions requires the knowledge of the in-cell protein concentrations. We determined the in-cell concentration of all mutants labeled with **GdI** from the DEER background decay. For frozen solutions with a homogenous distribution the slope of the background decay is a function of the spin concentration (see SI for details) (76, 56, 46). We generated a calibration curve from a plot of the slope of the background decays for all solutions C12A/A28C-**GdI** in buffer against their respective protein concentrations (Figure 4d) and this was used to determine local in-cell concentrations of delivered BIR1 (red symbols, Fig. 4c). Given the paramagnetic contributions of endogenous Mn^2+^, delineated protein concentrations clearly represented upper-limit estimates and should be corrected. Details of this procedure applied therein are described in SI and Fig. S20. Realistic local in-cell concentrations were in the range of 20-90 μM, 10–30% smaller than indicated by the primary data (Table S1, SI). Based on these results, we back-calculated expected λ’s assuming that dimer-monomer equilibria (and *K*_D_ values) are the same in buffer and in cells. By comparing calculated λ’s with experimental values, we noted large differences that clearly exceeded contributions by inherent experimental errors (Fig. 6). For these reasons, we were unable to attribute the significantly lower in-cell λ values to reduced intracellular BIR1 concentrations alone. This conclusion was further supported by the only marginally lower modulation depths of 30 μM BIR1 in cell lysates, compared to the 200 μM buffer samples (Fig. 3). Accordingly, we reasoned that low in-cell λ’s may reflect altered dimer-monomer equilibria in cells, with a significantly larger portion of monomeric BIR1 and a greater intracellular dimer-dissociation constant. These considerations stipulated that the BIR1 dimer may be subject to substantial destabilization in intact HeLa cells. Using the experimental in-cell λ values and the in-cell corrected protein concentrations for the five conjugates (WT BIR1, C12A/K19C, C12A/E22C, C12A/V24C and C12A/N28C), given in Table S1, we calculated the in-cell *K*_D_ for each of the measured BIR1-**GdI** conjugates under the assumption that *K*_D_ is independent of the labeling site and obtained *K*_D_= 126 ± 40 μM.

**Fig. 6.**
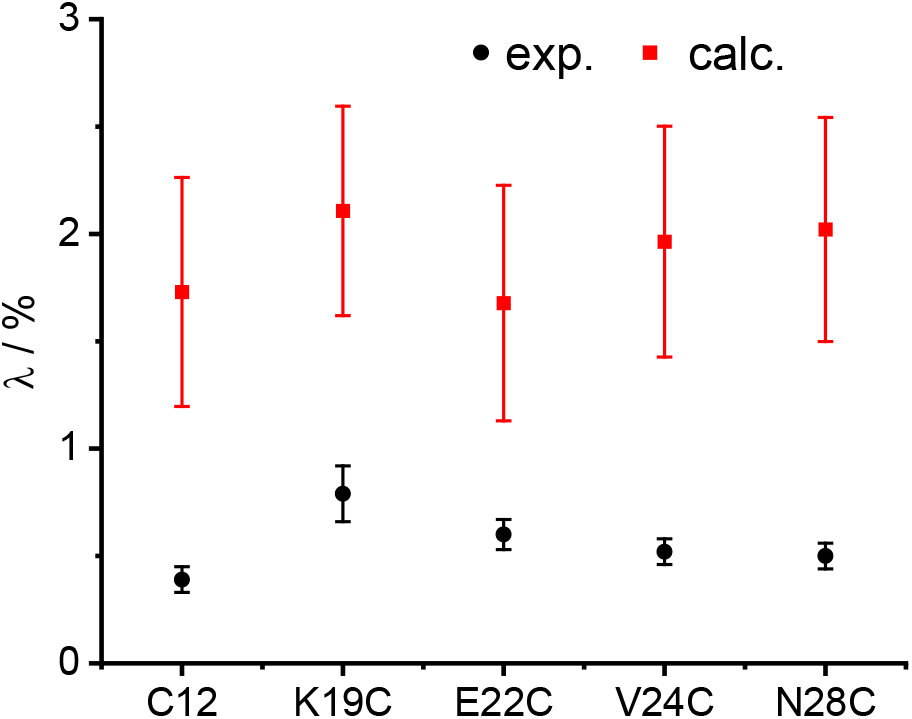
Comparison of the in-cell experimental λ values for **GdI** labeled BIR1 conjugates inside HeLa cells (black symbols) and in-cell expected λ values (red squares) calculated using the solution *K*_D_ value and corrected in-cell local concentrations determined from the solution calibration curve (Figure 4c).

To exclude the possibility that reduced in-cell modulation was a result of in-cell phosphorylation at residue S87 we carried out in-cell DEER measurements on the S87A mutant that cannot be phosphorylated. Specifically, we studies C12A/S87A/N28C–**GdI** and C12A/S87A/E22C–**GdI** (Fig. S20, SI) and both samples showed marginal or no modulations in the cell; that is, the amounts of dimers were below the detection limit. We determined the local in-cell concentrations of C12A/S87A/E22C–**GdI** and C12A/S87A/N28C-**GdI** from the calibration curve shown in Fig. 5d and obtained ~90 μM and ~160 μM, respectively. Considering a 25% reduction due to Mn^2+^ contributions, as explained above, this reduces to 67 and 120 μM. For such concentrations the DEER effect in solution was significant (Fig. 5), indicating that that also for the S87A mutant, the solution *K*_D_ was smaller than the in-cell *K*_D_. We, thus, conclude from these measurements that the decreased λ in cells is owing to the increase in *K*_D_ and does not originate from phosphorylation of BIR1 in HeLa cells after delivery.

### Co-solute effects on BIR1-dimer dissociation constants

To understand the nature of altered dimer dissociation constants in cells, we performed DEER experiments under artificially crowded *in vitro* conditions (77). In a first step, we employed Ficoll (300 g/L) as a crowding agent and measured C12A/S87A/N28C–**GdI** BIR1 modulation depths at various concentrations. At 100 μM of protein, we observed no modulations, which indicated that complete dimer dissociation had occurred (Fig. S21a, b). Addition of 150 g/L and 300 g/L of Ficoll to 150 μM BIR1 reduced modulation depths, suggesting progressive dimer destabilization and enhanced dissociation (Figure S21c, d). Interestingly, Ficoll effects appeared particularly pronounced at low BIR1 concentrations (<150 μM). In contrast to Ficoll, addition of BSA (300 g/L) and lysozyme (30 g/L) (exposure time of 10 min) as model agents for biological crowders(77) only marginally impacted BIR1-dimer stability (Fig. S22). We attempted to confirm Ficoll-mediated BIR1 destabilization effects by NMR spectroscopy. Unfortunately, we observed site-selctive line-broadening for 160 μM and 80 μM of BIR1 at increasing amounts of Ficoll (up to 300 g/L), without major chemical shift changes (Figure S23). NMR signals of flexible N- and C- terminal BIR1 residues remained visible whereas resonance peaks of the folded BIR1 domain progressively vanished. Thus, reduced rotational tumbling of BIR1 at high Ficoll concentrations prevented a detailed dimer-monomer analysis by NMR spectroscopy. Because the BIR1-BIR1 interface critically depends on a conserved salt bridge (67), we also tested how physiological salt concentrations (~0.2 M)(78, 79) affected dimer stability. We performed DEER measurements on 100 μM C12A/N28C-**GdI** BIR1 at 0.1-0.3 M of sodium chloride (Fig. S24). We clearly detected salt-dependent changes of modulation depths, which established that BIR1 dimers were salt sensitive and that weakening of electrostatic interactions at the dimer interface similarly resulted in BIR1 dissociation. In comparison to Ficoll, salt effects were not as prominent. Fig. 7 summarizes the effects of co-solutes on BIR1 modulation depths as proxies for respective dimer stabilities.

**Fig. 7.**
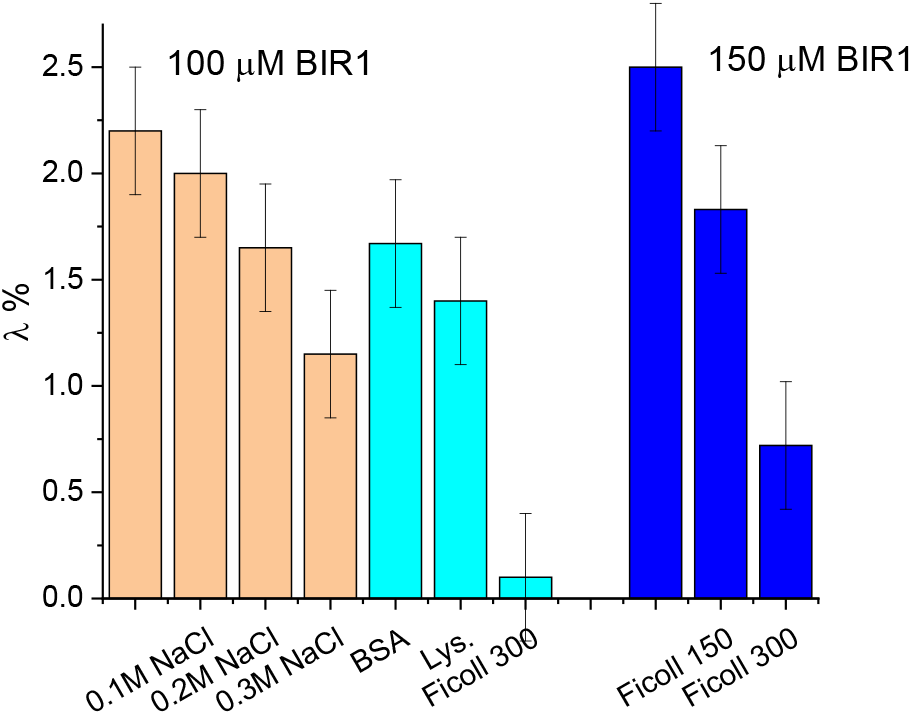
Co-solute effects on BIR1-dimer stability. Changes of C12A/N28C-**GdI** BIR1 modulation depths in the presence of salt (orange) and different macromolecular crowding agents (cyan and blue). Ficoll 150 and 300 correspond to mg/mL. Respective BIR1 concentrations are indicated.

We observed that the stability of the BIR1 dimer in dilute lysates was comparable to that in buffer. By contrast, BIR1 stability in lysed in-cell EPR samples was similar to the one measured in intact cells (Fig. 3b). We reasoned that this discrepancy may reflect differences in BIR1 to total cellular content concentrations ratios in the two types of lysate samples. For primary lysate samples, we employed 40,000 cells/μL that we cleared by centrifugation to sediment membranes and other insoluble components. Compared to intact HeLa cells, this corresponded to a ~10-fold dilution of soluble, cytoplasmic constituents (80). To these mixtures, we added 30 μM of spin-labeled BIR1, which resulted in a large excess of recombinant over endogenous proteins. Accordingly, dimer stability was comparable to pure-buffer conditions. By contrast, lysis of in-cell EPR samples produced crude solutions of low BIR1 concentrations against high amounts of cellular components. In turn, measured modulation depths and delineated *K*_D_’s were similar to intact cells.

## Discussion

In this study, we used W-band DEER distance measurements on the homo-dimeric BIR1 domain of XIAP, spin-labeled with **GdI** and **GdII**, to investigate dimer stability in HeLa cells, primary cell lysates and buffer. We found that (i) BIR1 dimers were destabilized in cells compared to dilute cell lysates and buffer. (ii) Dimer destabilization in buffer was observed in the presence of the artificial crowding agent Ficoll and at higher salt concentrations. (iii) N-terminal BIR1 segments spanning E22 experienced reduced conformational flexibility in intact cells. By contrast, flexible regions around C12 (disordered) and helical residues 26–31 (folded) maintained their respective structural states in all three environments.

First, we discuss changes in conformational space and local ordering within the disordered, N-terminal region in light of expected macromolecular crowding effects poised to limit conformational flexibility and lead to different levels of structural compaction (4, 6). Such behaviors are predicted by classic excluded volume theory (hard-sphere repulsion) and have been experimentally verified on a number of unstructured proteins (9, 41, 81). For example, compaction has been experimentally demonstrated by an artificial special FRET sensor, designed to probe conformational changes due to crowding (82). The sensor behavior was studied in solution in the presence of various crowding agents and in prokaryotic and eukaryotic cells (*E. Coli* and HEK293 cells) confirming the preference of more compact conformations as crowding increases. The behavior of structured proteins is however, more complex and depends on the protein. A recent in-cell NMR study on GB1 in eukaryotic cells reported a significant shift in the α-helix position compared to solution structure (28) presumably due to non-specific interactions of hydrophobic residues with cell components. In contrast, the structure of a truncated form of apoSOD1, was unaffected by the cellular environment (83). For BIR1 we did not observe an obvious overall compaction of the disordered part of BIR1 in cells. The C12 site exhibited a very broad distance distribution in all environments but small changes may be difficult to detect under these conditions. The N28C labeling site revealed a minor reduction, 0.2 nm, in the maximum of the distance distribution in the cell and addition of Ficoll led to a similar subtle shortening (see Fig. 22d) but these are too small to justify a conclusions regarding compaction due to crowding. In-cell distance distribution at E22C, closer to folded BIR1 parts, were somewhat narrower, with reduced intensity at the shorter distance range. This change reflects reduced dynamics potentially coupled to a local conformational change. To our surprise, we were unable to reproduce these E22 effects in the presence of Ficoll, BSA or salt, and the reduced flexibility was not detected in primary cell lysates. The in-cell changes are presumably caused by a combination of cellular crowding and soft interactions (84). We excluded that the change arises from binding of the TAB1 partner (69) based on stoichiometry considerations and the high intracellular amounts of delivered BIR1 (~ 50 μM), exceeding native TAB1 concentrations by at least one order of magnitude. Clearly, the most interesting finding of our study relates to the reduced stability of the BIR1 homodimer in intact HeLa cells. The estimated destabilization free energy determined from the dissociation constants amounts to ΔΔG^0^=1.24-1.31 Kcal/mol, for a temperature range of 260-273 K. A range is given owing to the uncertainty of the freezing temperature. In the following paragraphs, we discuss these results in the context of emerging concepts of how crowding and dimer shapes (85, 86), as well as surface charges (87–90), contribute to dimer cellular stabilities. According to excluded volume theory (91, 92), protein association is favored under conditions of macromolecular crowding with concomitant increases in complex stabilities (7, 85). These predictions are based on the notion that cellular proteins strive to adopt states of lowest possible volume occupancies. While this may lead to disordered protein compaction, it may similarly enhance protein association, especially when the association results in a volume decrease. In a rough approximation, this holds true for protein dimers that adopt spherical shapes, which represent ideal monomer ‘packing’. In turn, spherical shapes of associated proteins translate into enhanced intracellular stabilities. By the same token, protein dimers, the shapes of which deviate from spherical shapes are expected to display reduced stabilities in crowded environments and can even be destablized(85). Such a shape-dependent behavior was recently confirmed experimentally *in-vitro* for two engineered dimers of the *Streptococcal* protein G B1 domain (GB1) which have different eccentricity (86). In the presence of Ficoll and PEG, a domain-swapped GB1 dimer with a lower eccentricity was significantly stabilized, whereas for the side-by-side dimer of GB1 with a higher eccentricity the effect in Ficoll was marginal and destabilization was observed in PEG solutions. In light of these findings we looked into the shape of the BIR1 dimer. Here we used the crystal structure (Fig. 1a), which does not include the 1-19 N-terminal disordered tail. This is an approximation because the monomer/dimer sizes and shapes in solution with this tail may be somewhat different. NMR measurements on a monomeric mutant showed that its structure in solution is very similar to that of the monomer within the dimer (67) and therefore we used the structure of the monomer in the dimer crystal structure to describe the monomer. The BIR1 dimer is a side-by-side dimer, where the volume of the dimer (19959 Å^3^) is slightly higher than twice that of the monomer (9451 Å^3^) and the accessible surface area (ASA, 8554 Å^2^) is slightly lower than that of the two monomers (9552 Å^2^). The dimer-monomer difference, ΔASA_D->M_=982 Å^2^, is comparable to that of the side by side swapped dimer of GB1 (ΔASA_D->M_=1020 Å^2^)(86). The shape of the BIR1 dimer is far from being a sphere and is best described by a trapezoid (see Fig. S25), whereas the monomer is closer to a globular shape. Therefore, the BIR1-dimer destabilization in crowded Ficoll environments and intact cells seems to agree with the predicted lack of stabilization, though the large destabilization is still puzzling. Finally, we cannot exclude that the flexible N-terminal segments also adds to the reduced stability of dimeric BIR1.

Taking into account only excluded volume effects on in-cell dimer stability is, however, insufficient. Computer simulations and experimental work have shown that soft interactions as well as diffusion can modulate protein-protein interactions (10, 87, 93). Contributions can also come from interaction with cellular metabolites and other physiological, small-molecule compounds(94). Collectively, these factors may stabilize or destabilize associated proteins depending on specific solvation interactions that energetically disfavors/favors individual monomers, respectively. Accordingly, a unified behavior is not expected, as indeed reported by the few reports on in-cell association of proteins. The association rate constants of TEM1 β-lactamase binding to its protein inhibitor β-lactamase inhibitor protein (BLIP) in the cytoplasm of HeLa cells revealed only subtle differences in binding kinetics between *in vitro* and *in vivo* conditions (23). In contrast to the BIR1 behavior, the *K*_D_ of the AcGFP1/mCherry FRET pair in PBS buffer (*K*_D_ ~ 20 μM) was found to decrease by an order of magnitude in the cell (*K*_D_= 2.0 ± 0.5 μM) (24). This expected increase was attributed to crowding or quinary interactions. Similarly, cross linking experiments studying the dimer-monomer equilibrium of HSP27 in cells showed that while some dimers could be found in the cell, after homogenizing the cells in a dilute buffer all dimers dissociated. The difference was assigned to the reduced hydration in cell, which promotes charge-charge interaction (95). Finally, an opposite behavior; the propensity fPGK to self-aggregate was found to decrease gradually as cell volume decrease and crowding increase (94).

We also explored the electrostatic (and hydrophobic) surface properties of the BIR1 dimers and monomer that can affect charge-charge interactions shown to modulate protein-protein interactions (87). The electrostatic surface potential of the dimer and monomer of BIR1 are presented in Fig. S25. The overall charge of BIR1 (calculated pI=6.08) is negative and it is relevant to compare its *in-vitro* behavior to that of the side-by-side GB1 dimer. The latter was studied in cell lysate and in solution in the presence of macromolecular co-solutes that mimic attractive and repulsive interactions to highlight the importance of chemical interactions (77). The highest stabilization was obtained in *E. coli* lysate, which was similar to that observed by the addition of the negatively charged BSA, whereas positively charged lysozyme destabilized the dimer. As the GB1 dimer has an overall negative charge, the stabilization effect of BSA was assigned to charge induced repulsion and the destabilization by lysozyme was a results of the charge attraction. A similar explanation was given for the effect of the cell lysate, which was described as having a majority of negatively charged proteins at the pH used (77). The *in vitro* behavior BIR1 is different; neither BSA, nor lysozyme or cell lysate had significant effects on the BIR1 dimer stability.

Increasing the salt concentration in solution, to account for the in-cell ionic strength, did reduce the dimer stability. This is expected owing to screening effects that destabilize the salt bridge (96) holding the BIR1 dimer together (67). We note, however, that the effect of salt addition was not as strong as observed in the cell. The effect of Ficoll on ion pairs was recently investigated using a sensor designed to report on the ionic strength in the cell (96). Such a sensor with a helix-pair rich in arginine and aspartic acid residues, which comprise the same pair forming the salt bridge in the BIR1 dimer, it was found that Ficoll compressed the sensor and this was attributed to the expected crowding effects. Therefore, we find the possibility that Ficoll perturbs the salt bridge in BIR1 dimer to be unlikely. Accordingly, we attribute the destabilization by Ficoll to a combination of crowding as described above and some chemical interactions (97). Finally, we cannot exclude the possibility that interaction with small molecule components of the cell affect the dimer stability as recently demonstrated for self-self aggregation of fPGK (FRET-labeled phosphoglycerate kinase (94).

To conclude, we attribute decrease dimer stability in the cell to the higher in-cell ionic strength, excluded volume effects combined with the side-by-side structure of the dimer, and possibly also additional contributions from interactions with cell components. It is tempting to assign the latter to quinary interactions, the high concentration of the delivered protein argues against it. Finally, because interaction of the BIR1 dimer with TAB1 was reported as essential for activation of the NF-κB pathway (41), the higher in-cell *K*_D_ may have implications for BIR1 function as for realistic low physiological concentrations the monomer would be the dominating form. However, the in-cell *K*_D_ for XIAP may be different than that of the isolated BIR1.

## Conclusion

By high field DEER measurements on BIR1 (the first BIR domain of XIAP) labeled with rigid **GdI** and **GdII** spin labels we explored the effect of the cellular environment on BIR1 dimer stability and the conformation of its flexible N-terminal domain. We found that the BIR1 dimer is destabilized in HeLa cells, and its dissociation constant, *K*_D_, is higher than in buffer solution and in dilute cell lysates. A destabilization of the BIR1 dimer was achieved in solution by the addition of the crowding agent Ficoll, as well as by increasing the salt concentration. This reduced dimer stability is explained by a combination of excluded volume effects combined with the dimer shape, the higher in-cell ionic and potentially other interactions with cell constituents. In addition, we observed that the N-terminal region containing residue E22 of BIR1 acquired somewhat lower conformational freedom in cells than in solution or in cell lysate, but this could not be reproduced in-vitro.

The in-cell behavior of BIR1, which showed a seemingly unexpected destabilization of the dimers, is an important example of the complexity of the cell environment and the diversity of its effects on protein’s behavior. This, and the scarce literature reports on in-cell protein-protein associations show that the nature of the protein itself is an important factor and is responsible for the different behaviors observed. This highlights the crucial need for a wealth of in-cell experimental data that will allow identifying general trends and correlations and thus improve our understanding of protein’s behavior in cells. Finally, we showed that DEER with the appropriate spin labels is an effective method for exploring protein properties in the cell and is expected to add to the new data and insights needed.

## Methods

### Protein preparation

Recombinant BIR1 of XIAP was expressed and purified as described previously.(66) Detailed procedures can be found in <u>*SI Appendix*</u>.

### Site-specific labeling with lanthanide tags or a fluorescent tag

For labeling with BrPSPy-DO3MA-Ln or BrPSPy-DO3A-Gd, the reaction was complete and purified as reported in the literature.(52, 53) The labeling efficiency for BrPSPy-DO3A-Gd was 66%, as determined by comparison of the echo-detected electron paramagnetic resonance (EPR) intensity of the tag alone and that of the labeled protein. The labeling efficiency of BrPSPy-DO3MA-Gd exceeded 80%. For EPR measurements in the solution or with co-factors, the spin-labeled protein concentration was 200 μM (per monomer) in 7:3 (v/v) Tris-D_2_O/glycerol-*d*_8_, unless otherwise noted. Fluorescence labeling was done by mixing WT BIR1 with ATTO488-M (Sigma) and TCEP at pH 6.5. Detailed procedures can be found in <u>*SI Appendix*</u>.

#### In-cell protein delivery

Proteins were delivered to HeLa cells by means of a previously reported electroporation procedure.(42) Briefly, the cells were suspended in electroporation buffer containing 0.25 mM labeled BIR1, and electroporated by means of a Nucleofector 2b device (Lonza) with pulse program B28 for HeLa cells. The cells were incubated at 37 °C for 5 h, and then detached and washed twice to remove the non-internalized protein and dead cells, and incubated in phosphate buffer solution containing 7:3 (v/v) D_2_O/glycerol-*d*_8_. The cells were counted on a hemocytometer before loading into an EPR capillary. The EPR samples in the capillary contained 200,000–300,000 cells/μL. Lysates of cells that had been electroporated with BIR1 conjugates were prepared by repeated freeze–thaw cycles in cryosurgery after the cells were loaded into the capillary. Detailed procedures can be found in the <u>*SI Appendix*</u>.

#### Preparation of cell lysates

HeLa cell lysates were prepared as described previously (70) The detached cells was re-suspended in 1 volume equiv. of cell lysis buffer. The BIR1 conjugates (30 μM) were mixed with cell extract, and subjected to the electroporation program used for in-cell samples with incubation at 37 °C for 5 h. For EPR measurements, glycerol-*d*_8_ (30% of the total volume) was added. Finally the cell extract samples contained 40,000 cells/μL. Detailed procedures can be found in the <u>*SI Appendix*</u>.

#### NMR spectroscopy

All NMR experiments were performed on a Bruker Avance 600 MHz NMR spectrometer equipped with a QCI CryoProbe. Unless noted otherwise, the NMR spectra of BIR1 and its mutants were recorded for the ^15^N-labeled protein sample at a concentration of 0.15 mM, a pH of 6.5, and a temperature of 298 K. Detailed procedures can be found in the <u>*SI Appendix*</u>.

#### EPR spectroscopy

All EPR measurements were carried out at 10K on a home-built W-band spectrometer (94.9 GHz) (98, 99). Echo-detected EPR (ED-EPR) spectra were recorded using the Hahn echo sequence (π/2-τ-π-τ-echo). The echo decays were measured with the same sequence, setting the magnetic field to the maximum of the ED-EPR spectra and varying τ. DEER measurements were recorded using a modified four-pulse DEER sequence with chirp pulse(s)(100–102).The DEER data were analyzed using the DeerAnalysis 2018 program with Tikhonov regularization (103). All experimental parameters are given in the <u>*SI Appendix*</u>.

#### Immunofluorescence microscopy

BIR1–ATTO488 was delivered to cells by means of the electroporation procedure used for Gd^3+^ labeled protein and the cells were recovered at 37 °C for 4 h. Cells were fixed with 4% paraformaldehyde. After washes, the cells were mounted and then imaged with an Olympus microscope X83 with a 60× oil objective. A 488 nm laser (Toptica, 100 mW) was used for fluorescence excitation, and a green LED was used for brightfield imaging. All experimental details are given in the <u>*SI Appendix*</u>.

The protein volume and surface area were obtained using the Volume Area Dihedral Angle Reporter (VADAR)(104) and the BIR1 dimer structure PDB code 2QRA (68). The protein pI and its total charge we obtained using the ProtParam tool of the ExPaSy software(105). Here one full monomer chain including the first 19 amino acid, not included in the crystal structure, was used.

## Acknowledgements

This work was supported by National Natural Science Foundation of China (NSFC)-Israel Science Foundation (ISF) grant (grant number 21761142004 and 2484/17,) to X.C.S. and D. G, Major National Scientific Research Project in China (2016YFA0501202), the Natural Science Foundation of China (21673122 and 21473095) to X. C. S. This research was made possible in part by the historic generosity of the Harold Perlman Family (D. G.). D. G. holds the Erich Klieger Professorial Chair in Chemical Physics. We thank Yasmin Ben Yishay for her help with sample preparation and Gideon Schreiber for very helpful discussions. Angeliki Giannoulis and Phil Selenko for critical reading of the manuscript.

## References

1. R. D. Cohen, G. J. Pielak, A cell is more than the sum of its (dilute) parts: A brief history of quinary structure. Protein Sci. 26, 403–413 (2017).

2. R. J. Ellis, Macromolecular crowding: an important but neglected aspect of the intracellular environment. Curr. Opin. Struct. Biol. 11, 114–119 (2001).

3. D. I. Freedberg, P. Selenko, Live cell NMR. Ann. Rev. Biophys. 43, 171–192 (2014).

4. D. Guin, M. Gruebele, Weak chemical interactions that drive proteine evolution: Crowding, sticking, and quinary structure in folding and function. Chem. Rev. 119, 10691–10717 (2019).

5. J. M. Plitzko, B. Schuler, P. Selenko, Structural Biology outside the box-inside the cell. Curr. Opin. Struct. Biol. 46, 110–121 (2017).

6. G. Rivas, A. P. Minton, Toward an understanding of biochemical equilibria within living cells. Biophys. Rev. 10, 241–253 (2018).

7. H.-X. Zhou, G. Rivas, A. P. Minton, Macromolecular crowding and confinement: Biochemical, biophysical, and potential physiological consequences. Ann. Revi. Biophys. 37, 375–397 (2008).

8. E. H. McConkey, Molecular evolution, intracellular organization, and the quinary structure of proteins. Proc. Natl. Acad. Sci. U S A 79, 3236–3240 (1982).

9. I. Yu et al., Biomolecular interactions modulate macromolecular structure and dynamics in atomistic model of a bacterial cytoplasm. eLife 5, e19274 (2016).

10. S. R. McGuffee, A. H. Elcock, Diffusion, crowding & protein stability in a dynamic molecular model of the bacterial cytoplasm. PLoS Comput. Biol. 6, e1000694 (2010).

11. G. Rivas, A. P. Minton, Macromolecular crowding in vitro, In vivo, and in between. Trends Biochem. Sci. 41, 970–981 (2016).

12. S. Qin, H. X. Zhou, Protein folding, binding, and droplet formation in cell-like conditions. Curr. Opin. Struct. Biol. 43, 28–37 (2017).

13. M. S. Cheung, A. G. Gasic, Towards developing principles of protein folding and dynamics in the cell. Phys. Biol. 15, 063001 (2018).

14. A. P. Minton, The influence of macromolecular crowding and macromolecular confinement on biochemical reactions in physiological media. J. Biol. Chem. 276, 10577–10580 (2001).

15. R. D. Cohen, G. J. Pielak, Electrostatic Contributions to qrotein quinary structure. J. Am. Chem. Soc. 138, 13139–13142 (2016).

16. R. D. Cohen, A. J. Guseman, G. J. Pielak, Intracellular pH modulates quinary structure. Protein Sci. 24, 1748–1755 (2015).

17. J. Danielsson et al., Thermodynamics of protein destabilization in live cells. Proc. Natl. Acad. Sci. U S A 112, 12402–12407 (2015).

18. W. B. Monteith, R. D. Cohen, A. E. Smith, E. Guzman-Cisneros, G. J. Pielak, Quinary structure modulates protein stability in cells. Proc. Natl. Acad. Sci. U S A 112, 1739–1742 (2015).

19. L. A. Benton, A. E. Smith, G. B. Young, G. J. Pielak, Unexpected effects of macromolecular crowding on protein stability. Biochemistry 51, 9773–9775 (2012).

20. D. Gnutt, M. Gao, O. Brylski, M. Heyden, S. Ebbinghaus, Excluded-volume effects in living cells. Angew. Chem. Int. Ed. Engl. 54, 2548–2551 (2015).

21. A. Dalaloyan et al., Tracking conformational changes in calmodulin in vitro, in cell extract, and in cells by electron paramagnetic resonance distance measurements. ChemPhysChem 20, 1860–1868 (2019).

22. F. X. Theillet et al., Physicochemical properties of cells and their effects on intrinsically disordered proteins (IDPs). Chem. Rev. 114, 6661–6714 (2014).

23. Y. Phillip, V. Kiss, G. Schreiber, Protein-binding dynamics imaged in a living cell. Proc. Natl. Acad. Sci. U S A 109, 1461–1466 (2012).

24. S. Sukenik, P. Ren, M. Gruebele, Weak protein-protein interactions in live cells are quantified by cell-volume modulation. Proc. Natl. Acad. Sci. U S A 114, 6776–6781 (2017).

25. A. Margineanu et al., Screening for protein-protein interactions using Forster resonance energy transfer (FRET) and fluorescence lifetime imaging microscopy (FLIM). Sci. Rep. 6, 28186 (2016).

26. K. Kwapiszewska et al., Determination of oligomerization state of Drp1 protein in living cells at nanomolar concentrations. Sci. Rep. 9, 5906 (2019).

27. D. Sakakibara et al., Protein structure determination in living cells by in-cell NMR spectroscopy. Nature 458, 102–105 (2009).

28. T. Tanaka et al., High-resolution protein 3D structure determination in living eukaryotic cells. Angew. Chem. Int. Ed. Engl. 58, 7284–7288 (2019).

29. T. L. X. Song, J. Chen, J. Wang, L. Yao, Characterization of residue specific protein folding and unfolding dynamics in cells. J. Am. Chem. Soc. 141, 11363–11366 (2019).

30. T. Muntener, D. Haussinger, P. Selenko, F. X. Theillet, In-cell protein structures from 2D NMR experiments. J. Phys. Chem. Lett. 7, 2821–2825 (2016).

31. B. B. Pan et al., 3D structure determination of a protein in living cells using paramagnetic NMR spectroscopy. Chem. Commun. 52, 10237–10240 (2016).

32. E. Luchinat, L. Banci, In-cell NMR in human cells: direct protein expression allows structural studies of protein folding and maturation. Acc. Chem. Res. 51, 1550–1557 (2018).

33. K. Inomata et al., High-resolution multi-dimensional NMR spectroscopy of proteins in human cells. Nature 458, 106–109 (2009).

34. S. Ebbinghaus, A. Dhar, J. D. McDonald, M. Gruebele, Protein folding stability and dynamics imaged in a living cell. Nat. Methods 7, 319–323 (2010).

35. M. Guo, Y. Xu, M. Gruebele, Temperature dependence of protein folding kinetics in living cells. Proc. Natl. Acad. Sci. U S A 109, 17863–17867 (2012).

36. A. Dhar et al., Protein stability and folding kinetics in the nucleus and endoplasmic reticulum of eucaryotic cells. Biophys. J. 101, 421–430 (2011).

37. I. Guzman, H. Gelman, J. Tai, M. Gruebele, The extracellular protein VlsE is destabilized inside cells. J. Mol. Biol. 426, 11–20 (2014).

38. W. B. Monteith, G. J. Pielak, Residue level quantification of protein stability in living cells. Proceedings of the National Academy of Sciences of the United States of America 111, 11335–11340 (2014).

39. A. P. Schlesinger, Y. Wang, X. Tadeo, O. Millet, G. J. Pielak, Macromolecular crowding fails to fold a globular protein in cells. J. Am. Chem. Soc. 133, 8082–8085 (2011).

40. D. Gnutt et al., Stability effect of quinary interactions reversed by single point mutations. J. Am. Chem. Soc. 141, 4660–4669 (2019).

41. M. M. Dedmon, C. N. Patel, G. B. Young, G. J. Pielak, FlgM gains structure in living cells. Proc. Natl. Acad. Sci. USA 99, 12681 (2002).

42. F. X. Theillet et al., Structural disorder of monomeric alpha-synuclein persists in mammalian cells. Nature 530, 45–50 (2016).

43. I. Konig et al., Single-molecule spectroscopy of protein conformational dynamics in live eukaryotic cells. Nat. Methods 12, 773–779 (2015).

44. M. Azarkh, O. Okle, P. Eyring, D. R. Dietrich, M. Drescher, Evaluation of spin labels for in-cell EPR by analysis of nitroxide reduction in cell extract of Xenopus laevis oocytes. J. Magn. Reson. 212, 450–454 (2011).

45. R. Igarashi et al., Distance determination in proteins inside Xenopus laevis oocytes by double electron-electron resonance experiments. J. Am. Chem. Soc. 132, 8228–8229 (2010).

46. J. J. Jassoy et al., Versatile trityl spin labels for nanometer distance measurements on biomolecules in vitro and within cells. Angew. Chem. Int. Ed. Engl. 56, 177–181 (2017).

47. G. Karthikeyan et al., A bioresistant nitroxide spin label for in-cell EPR spectroscopy: In vitro and in oocytes protein structural dynamics studies. Angew. Chem. Int. Ed. Engl. 57, 1366–1370 (2018).

48. I. Krstic et al., Long-range distance measurements on nucleic acids in cells by pulsed EPR spectroscopy. Angew. Chem. Int. Ed. Engl. 50, 5070–5074 (2011).

49. A. Martorana et al., Probing protein conformation in cells by EPR distance measurements using Gd3+ spin labeling. J. Am. Chem. Soc. 136, 13458–13465 (2014).

50. F. C. Mascali, H. Y. Ching, R. M. Rasia, S. Un, L. C. Tabares, Using genetically encodable self-assembling Gd(III) spin labels to make in-cell nanometric distance measurements. Angew. Chem. Int. Ed. Engl. 55, 11041–11043 (2016).

51. M. Qi, A. Gross, G. Jeschke, A. Godt, M. Drescher, Gd(III)-PyMTA label is suitable for in-cell EPR. J. Am. Chem. Soc. 136, 15366–15378 (2014).

52. Y. Yang et al., High sensitivity in-cell EPR distance measurements on proteins using an optimized Gd(III) spin label. J. Phys. Chem. Lett. 9, 6119–6123 (2018).

53. Y. Yang et al., A reactive, rigid Gd(III) labeling tag for in-cell EPR distance measurements in proteins. Angew. Chem. Int. Ed. Engl. 56, 2914–2918 (2017).

54. Y. Yang, F. Yang, X. Y. Li, X. C. Su, D. Goldfarb, In-cell EPR distance measurements on ubiquitin labeled with a rigid PyMTA-Gd(III) tag. J. Phys. Chem. B 123, 1050–1059 (2019).

55. M. Azarkh et al., Long-range distance determination in a DNA model system inside Xenopus laevis oocytes by in-cell spin-label EPR. ChemBioChem 12, 1992–1995 (2011).

56. G. Jeschke, Y. Polyhach, Distance measurements on spin-labelled biomacromolecules by pulsed electron paramagnetic resonance. Phys. Chem. Chem. Phys. 9, 1895–1910 (2007).

57. E. R. Georgieva, P. P. Borbat, H. D. Norman, J. H. Freed, Mechanism of influenza A M2 transmembrane domain assembly in lipid membranes. Scientific reports 5, 11757 (2015).

58. J. Schredelseker et al., High resolution structure and double electron-electron resonance of the zebrafish voltage-dependent anion channel 2 reveal an oligomeric population. J. Biol. Chem. 289, 12566–12577 (2014).

59. S. Fulda, D. Vucic, Targeting IAP proteins for therapeutic intervention in cancer. Nat. Rev. Drug Discov. 11, 109–124 (2012).

60. A. D. Schimmer, S. Dalili, R. A. Batey, S. J. Riedl, Targeting XIAP for the treatment of malignancy. Cell Death Differ. 13, 179–188 (2006).

61. M. Gyrd-Hansen, P. Meier, IAPs: from caspase inhibitors to modulators of NF-kappaB, inflammation and cancer. Nat. Rev. Cancer 10, 561–574 (2010).

62. S. Galban, C. S. Duckett, XIAP as a ubiquitin ligase in cellular signaling. Cell Death Differ. 17, 54–60 (2010).

63. Y. Huang et al., Structural basis of caspase inhibition by XIAP: differential roles of the linker versus the BIR domain. Cell 104, 781–790 (2001).

64. S. J. Riedl et al., Structural basis for the inhibition of caspase-3 by XIAP. Cell 104, 791–800 (2001).

65. C. Sun et al., NMR structure and mutagenesis of the inhibitor-of-apoptosis protein XIAP. Nature 401, 818–822 (1999).

66. G. Wu et al., Structural basis of IAP recognition by Smac/DIABLO. Nature 408, 1008–1012 (2000).

67. M. M. Hou et al., Solution structure and interaction with copper in vitro and in living cells of the first BIR domain of XIAP. Sci. Rep. 7, 16630 (2017).

68. S. C. Lin, Y. Huang, Y. C. Lo, M. Lu, H. Wu, Crystal structure of the BIR1 domain of XIAP in two crystal forms. J. Mol. Biol. 372, 847–854 (2007).

69. M. Lu et al., XIAP induces NF-kappaB activation via the BIR1/TAB1 interaction and BIR1 dimerization. Mol. Cell 26, 689–702 (2007).

70. F. X. Theillet et al., Site-specific NMR mapping and time-resolved monitoring of serine and threonine phosphorylation in reconstituted kinase reactions and mammalian cell extracts. Nat. Protoc. 8, 1416–1432 (2013).

71. G. Hagelueken, R. Ward, J. H. Naismith, O. Schiemann, MtsslWizard: In silico spin-labeling and generation of distance distributions in PyMOL. Appl. Magn. Reson. 42, 377–391 (2012).

72. K. Shiraki, K. Nishikawa, Y. Goto, Trifluoroethanol-induced stabilization of the alpha-helical structure of beta-lactoglobulin: implication for non-hierarchical protein folding. J. Mol. Biol. 245, 180–194 (1995).

73. A. Giannoulis et al., Nitroxide-nitroxide and nitroxide-metal distance measurements in transition metal complexes with two or three paramagnetic centres give access to thermodynamic and kinetic stabilities. Phys. Chem. Chem. Phys. 20, 11196–11205 (2018).

74. K. Kato et al., Protein kinase C stabilizes X-linked inhibitor of apoptosis protein (XIAP) through phosphorylation at Ser87 to suppress apoptotic cell death. Psychogeriatrics 11, 90–97 (2011).

75. J. L. Wort et al., Sub-micromolar pulse dipolar EPR spectroscopy reveals increasing Cu(II)-labelling of double-histidine motifs with lower temperature. Angew. Chem. Int. Ed. 58, 11681–11685 (2019).

76. G. Jeschke, A. Koch, U. Jonas, A. Godt, Direct conversion of EPR dipolar time evolution data to distance distributions. J. Magn. Reson. 155, 72–82 (2002).

77. A. J. Guseman, G. J. Pielak, Cosolute and crowding effects on a side-by-side protein dimer. Biochemistry 56, 971–976 (2017).

78. G. K. J. Kort, The Na^+^, K^+^, 2Cl^−^-cotransport system in HeLa cells and HeLa cell mutants exhibiting an altered efflux pathway,. J. Cellular Physiol. 141, 181–190 (1989).

79. M. C. Jewett, J. R. Swartz, Substrate replenishment extends protein synthesis with an in vitro translation system designed to mimic the cytoplasm. Biotechnol. Bioeng. 87, 465–472 (2004).

80. L. K. Zhao, C. D.; Song, J.; Piwnica-Worms, D.; Ackerman, J. J.; Neil, J.J., Intracellular water-specific MR of microbead-adherent cells: the HeLa cell intracellular water exchange lifetime. NMR Biomed. 21, 159–164 (2008).

81. J. Hong, L. M. Gierasch, Macromolecular crowding remodels the energy landscape of a protein by favoring a more compact unfolded state. J. Am. Chem. Soc. 132, 10445–10452 (2010).

82. A. J. Boersma, I. S. Zuhorn, B. Poolman, A sensor for quantification of macromolecular crowding in living cells. Nat. Methods 12, 227–229 (2015).

83. J. Danielsson et al., Pruning the ALS-associated protein SOD1 for in-cell NMR. J. Am. Chem. Soc. 135, 10266–10269 (2013).

84. M. Sarkar, C. Li, G. J. Pielak, Soft interactions and crowding. Biophysical reviews 5, 187–194 (2013).

85. O. G. Berg, The influence of macromolecular crowding on thermodynamic activity: solubility and dimerization constants for spherical and dumbbell-shaped molecules in a hard-sphere mixture. Biopolymers 30, 1027–1037 (1990).

86. A. J. Guseman, G. M. Perez Goncalves, S. L. Speer, G. B. Young, G. J. Pielak, Protein shape modulates crowding effects. Proc. Natl. Acad. Sci. U S A 115, 10965–10970 (2018).

87. A. J. Guseman, S. L. Speer, G. M. Perez Goncalves, G. J. Pielak, Surface charge modulates protein-protein interactions in physiologically relevant environments. Biochemistry 57, 1681–1684 (2018).

88. X. Mu et al., Physicochemical code for quinary protein interactions in Escherichia coli. Proc. Natt. Acad. Sci. USA 114, E4556 (2017).

89. J. Danielsson, F. Yang, M. Oliveberg, “In-cell NMR: Spectral optimization and population adjustment by rationalprotein engineering” in In-cell NMR spectroscopy: From molecular sciences to cell biology, Y. Ito, V. Dötsch, M. Shirakawa, Eds. (RSC, 2019), 10.1039/9781788013079-00171 chap. 11, pp. 171–187.

90. A.J. Guseman, G. J. Pielak, “Protein Stability and Weak Intracellular Interactions” in In-cell NMR spectroscopy: From molecular sciences to cell biolog, Y. Ito, V. Dötsch, M. Shirakawa, Eds. (RSC, 2019), 10.1039/9781788013079-00188 chap. 12, pp. 188–206

91. A. P. Minton, The effect of volume occupancy upon the thermodynamic activity of proteins: some biochemical consequences. Mol. Cell. Biochem. 55, 119–140 (1983).

92. A. P. Minton, How can biochemical reactions within cells differ from those in test tubes? J. Cell Sci. 119, 2863–2869 (2006).

93. Y. Phillip, G. Schreiber, Formation of protein complexes in crowded environments--from in vitro to in vivo. FEBS Lett. 587, 1046–1052 (2013).

94. S. Sukenik, M. Salam, Y. Wang, M. Gruebele, In-cell titration of small solutes controls protein stability and aggregation. J. Am. Chem. Soc. 140, 10497–10503 (2018).

95. S. B. Sato, M. Sugiura, T. Kurihara, Dimer-monomer equilibrium of human HSP27 is influenced by the in-cell macromolecular crowding environment and is controlled by fatty acids and heat. Biochim. Biophys. Acta 1866, 692–701 (2018).

96. B. Liu, B. Poolman, A. J. Boersma, Ionic strength sensing in living cells. ACS Chem. Biol. 12, 2510–2514 (2017).

97. L. A. Benton, A. E. Smith, G. B. Young, G. J. Pielak, Unexpected effects of macromolecular crowding on protein stability. Biochemistry 51, 9773–9775 (2012).

98. D. Goldfarb et al., HYSCORE and DEER with an upgraded 95 GHz pulse EPR spectrometer. J. Magn. Reson. 194, 8–15 (2008).

99. F. Mentink-Vigier et al., Increasing sensitivity of pulse EPR experiments using echo train detection schemes. J. Magn. Reson. 236, 117–125 (2013).

100. T. Bahrenberg et al., Improved sensitivity for W-band Gd(III)-Gd(III) and nitroxide-nitroxide DEER measurements with shaped pulses. J. Magn. Reson. 283, 1–13 (2017).

101. Y. Y. Bahrenberg T., Goldfarb D., Feintuch A., rDEER: A modified DEER sequence for distance measurements using shaped pulses. Magnetochemistry 5, 20 (2019).

102. A. Doll et al., Gd(III)-Gd(III) distance measurements with chirp pump pulses. J. Magn. Reson. 259, 153–162 (2015).

103. G. Jeschke et al., DeerAnalysis2006—a comprehensive software package for analyzing pulsed ELDOR data. Appl. Magn. Reson. 30, 473–498 (2006).

104. https://academic.oup.com/nar/article/31/13/3316/2904187 (last access March 23, 2020).

105. https://web.expasy.org/protparam/ (last access 22.3.2020).

