## Supplementary information for "In-cell destabilization of a homo-dimeric protein complex detected by DEER spectroscopy"

#### Contents

**1. Experimental** ........ S2

*Protein preparation* ........ S2

*Site-specific labeling of BIR1 with lanthanide tags or a fluorescent tag* ........ S2

*In-cell protein delivery* ........ S3

*Preparation of cell lysates* ........ S3

*NMR spectroscopy* ........ S3

*EPR spectroscopy* ........ S3

*Immunofluorescence microscopy* ........ S4

**2. Mass-spectrometry data** ........ S5

**3. NMR measurements** ........ S6

**4. DEER data of monomeric BIR1** ........ S8

**5. EPR and echo-decays** ........ S9

**6. Primary DEER data and analysis for BIR1 mutants** ........ S10

**7. Primary DEER data and analysis for E22C and N28C in the presence of co-solutes** ........ S13

**8. Primary DEER data and analysis for K19C and V24C** ........ S15

**9. Determination of dissociation constants from DEER data** ........ S16

**10. Monomer-dimer equilibrium observed by NMR** ........ S17

**11. SEC results on WT BIR1, S87A and S87E mutants** ........ S18

**12. NMR relaxation data on WT BIR1, D71N/R72E, S87A and S87E mutants** ........ S19

**13. Determination of in-cell concentrations** ........ S19

**14. DEER results on S87A/C12A/E22C-GdI and S87A/C12A/N28C-GdI** ........ S22

**15. Effect of co-solutes on modulation depth** ........ S23

**16. Electrostatic potential surfaces** ........ S25

### 1. Experimental

#### *Protein preparation*

Recombinant wild-type (WT) BIR1 (residues 1–105) of XIAP and 13 mutants (C12A, C12A/K19C, C12A/V24C, C12A/E22C, C12A/N28C, D71N/R72E, C12A/E22C/D71N/R72E, C12A/N28C/D71N/R72E, S87E, C12A/E22C/S87E, S87A, C12A/E22C/S87A and C12A/N28C/S87A) were cloned into the PET3a vector. Unlabeled and $^{15}$N-labeled proteins were expressed in *E. coli* in LB medium and M9 medium with induction by isopropyl β-D-1-thiogalactopyranoside, respectively, and the resulting proteins were purified on a diethylaminoethyl column and then subjected to Superdex75 gel filtration, as described previously (1). In this way, approximately 18 mg of $^{15}$N-labeled BIR1 and 25 mg of unlabeled BIR1 were obtained from 250 mL of M9 and 1 L of LB media, respectively.

#### *Site-specific labeling of BIR1 with lanthanide tags or a fluorescent tag*

The lanthanide spin labels were synthesized as described previously (2,3). Then 1 mL of 0.3 mM WT BIR1 or a mutant in 20 mM tris(hydroxymethyl)aminomethane (Tris) buffer was mixed with 6 equiv (with respect to one free cysteine) of BrPSPy-DO3M(R)A-Ln (Ln = $Gd^{3+}$, $Dy^{3+}$, $Tm^{3+}$, or $Y^{3+}$) where R indicates that the carbon atom in the arm of 1,4,7,10-tetraazacyclododecane (cyclen) ring was in the *R* configuration, or BrPSPy- DO3A-Gd (in the form of 50 mM aqueous stock solutions) and 0.1 mM tris(2-carboxyethyl)phosphine. We refer to BrPSPy-DO3A-Gd as **GdI** and to BrPSPy- DO3MA-Gd as **GdII**. The pH of the resulting solution was adjusted to approximately 8 with 0.5 M NaOH, and the solution was then incubated at room temperature with monitoring by matrix-assisted laser desorption/ionization time-of-flight mass spectrometry (Fig. S1). Typically, the reaction was complete within 5–8 h depending on the reactivity of the cysteine at the ligation site. Free lanthanide tag was then removed with a short PD-10 desalting column. The overall yield of spin-labeled protein was about 70%, as indicated by UV absorption at 280 nm. The labeling efficiency for **GdI** was 66%, as determined by comparison of the echo-detected electron paramagnetic resonance (EPR) intensity of the tag alone and that of the labeled protein. The labeling efficiency of **GdII** exceeded 80%.

For solution EPR measurements, the spin-labeled proteins were dissolved in 7:3 (v/v) Tris-$D_2O$ (pD 7.2)/glycerol-$d_8$. Samples containing Ficoll 400 (Sigma-Aldrich, 150 or 300 g/L), chicken egg white lysozyme (Sigma-Aldrich, 300 g/L), or bovine serum albumin (BSA; Sigma-Aldrich, 300 g/L) as co-solutes were prepared in 7:3 (v/v) Tris-$D_2O$ (pD 7.2)/glycerol-$d_8$. Unless otherwise noted, the protein concentration was 200 μM (per monomer) for all DEER experiments in 20 mM Tris solution.

For fluorescence labeling, 1 mM WT BIR1 (0.1 mL) was mixed with 3.0 mM ATTO488-maleimide (ATTO488-M, Sigma-Aldrich), and 1.0 mM tris(2-carboxyethyl)phosphine in 20 mM (4-morpholineethanesulfonic acid) (MES) buffer at pH 6.5. The mixture was incubated at room temperature for 4 h. Unreacted fluorescent label was removed by means of size exclusion chromatography (HiLoad 16/60 Superdex 30).

***In-cell protein delivery***

Proteins were delivered to HeLa cells by means of a previously reported electroporation procedure (4). Briefly, the cells were suspended in 100 μL of PBS electroporation buffer (100 mM sodium phosphate, 5 mM KCl, 15 mM $MgCl_2$, 15 mM 4-(2-hydroxyethyl)piperazine-1-ethanesulfonic acid (HEPES), 2 mM ATP, 2 mM reduced glutathione, pH 7.4) containing 0.25 mM labeled BIR1. The suspension was then transferred to a 2 mm cuvette and electroporated by means of a Nucleofector 2b device (Lonza) with pulse program B28 for HeLa cells, as described by the manufacturer. The cells were then transferred to collagen-treated dishes, incubated at 37 °C for 5 h, and detached from the culture dishes with trypsin. The detached cells were centrifuged at 1000*g* for 5 min at 25 °C, and the pellet was washed twice in PBS-$D_2O$ buffer to remove the noninternalized protein and dead cells and then incubated for 10 min in PBS buffer containing 7:3 (v/v) $D_2O$/glycerol-$d_8$. The cells were counted on a hemocytometer and then loaded into an EPR capillary and centrifuged at 1500*g* for 30 min. The EPR samples in the capillary contained 200,000–300,000 cells/μL. Finally, the capillary was slowly frozen in an isopropanol rack at −80 °C. Lysates of cells that had been electroporated with BIR1 conjugates were prepared by repeated freeze–thaw cycles in cryosurgery after the cells were loaded into the capillary.

***Preparation of cell lysates***

HeLa cell lysates were prepared as described previously (5). When the cells reached 60–70% confluency, they were detached from the culture dishes with trypsin/EDTA (0.05%/0.02%), washed with ice-cold PBS-$D_2O$ (100 mM, pD 7.2), and sedimented by centrifugation at 1300*g* for 5 min. The cell pellet was resuspended in 1 volume equiv of cell lysis buffer (PBS-$D_2O$, supplemented with 150 mM NaCl, 1% NP-40, protease inhibitor cocktail III [Calbiochem], pD 7.2), and kept on ice for 15 min. The cell extract was snap-frozen and stored in liquid nitrogen until use. The BIR1 conjugates (30 μM) were mixed with 100 μL of cell extract, and the mixture was subjected to the electroporation program used for in-cell samples with incubation at 37 °C for 5 h. For EPR measurements, glycerol-$d_8$ (30% of the total volume) was added before the sample was loaded into an EPR capillary. The cell extract contained 40,000 cells/μL.

***NMR spectroscopy***

All NMR experiments were performed on a Bruker Avance 600 MHz NMR spectrometer equipped with a QCI CryoProbe. Unless noted otherwise, the NMR spectra of BIR1 and its mutants were recorded in 20 mM 2, 2-bis(hydroxyethyl)-(iminotris)-(hydroxymethyl)-methane (Bis-Tris) buffer for the $^{15}$N-labeled protein sample at a concentration of 0.15 mM, a pH of 6.5, and a temperature of 298 K.

***EPR spectroscopy***

All EPR measurements were carried out at 10K on a home-built W-band spectrometer (94.9 GHz) (6,7). W-band echo-detected EPR (ED-EPR) spectra were recorded by using the Hahn echo sequence (π/2-τ-π-τ-echo) with π/2 and π pulse durations of 15 and 30 ns, respectively, with τ = 550 ns, a repetition time of 1 ms, and sweeping of the magnetic field. We measured echo decays by means of the same sequence, setting the magnetic field to the maximum of the ED-EPR spectra and varying $\tau$.

DEER measurements were recorded by using a modified four-pulse DEER sequence (8). The observer pulses were set to the maximum of the EPR spectrum, and the $\pi/2$ and $\pi$ pulse durations were 15 and 30 ns, respectively. The pump pulse was set on both sides of the maximum of the central transition with a frequency offset of 100 MHz. The bandwidth of the chirp pump pulse was 300 MHz, and the duration was 96 ns. The second delay time was 5 μs, the step of *t* was 40 ns, and the repetition time was 800 μs. Standard four-pulse DEER sequences (9) were also used in this work, where indicated. The pump pulse duration was 15 ns, and the observer pulse durations were 15 and 30 ns respectively. The frequency difference between the pump and observer pulses was 100 MHz, with the pump pulse set to the maximum of the EPR spectrum. The first delay time was 400 ns, the step of *t* was 50 ns, and the repetition time was 800 μs. The accumulation time ranged from 4 to 8 h for measurements in solution and in lysates and from 10 to 24 h for in-cell measurements.

The DEER data were analyzed by using the DeerAnalysis 2018 program (10). Distance distributions were obtained by using Tikhonov regularization. Background was fitted with a homogenous three dimensional distribution, except when noted otherwise on the figure caption. Data were validated by varying the background start range from 15% to 70% of the primary time window in 11 trials.

***Immunofluorescence microscopy***

For immunofluorescence imaging, BIR1–ATTO488 was delivered to cells by means of the electroporation procedure used for $Gd^{3+}$ labeled protein. The cells were recovered in the incubator on collagen-treated 25 mm cover slips for 4 h. Cells were rinsed three times in PBS and then fixed in PBS containing 4% paraformaldehyde for 15 min. After being washed three times with PBS, the cells on the coverslips were mounted with a drop of PBS and sealed with nail polish. Confocal images were obtained with an automated Olympus microscope X83 with a 60× oil objective (Olympus, plan apo, 1.42 numerical aperture) coupled to a spinning disk confocal scanner (Yokogawa W1). A 488 nm laser (Toptica, 100 mW) was used for fluorescence excitation, and a green LED was used for brightfield imaging. The emission filter-sets used for brightfield and fluorescence images were identical (520/28, Chroma). Images were recorded with a Hamamatsu Flash4 camera.

### 2. Mass-spectrometry data

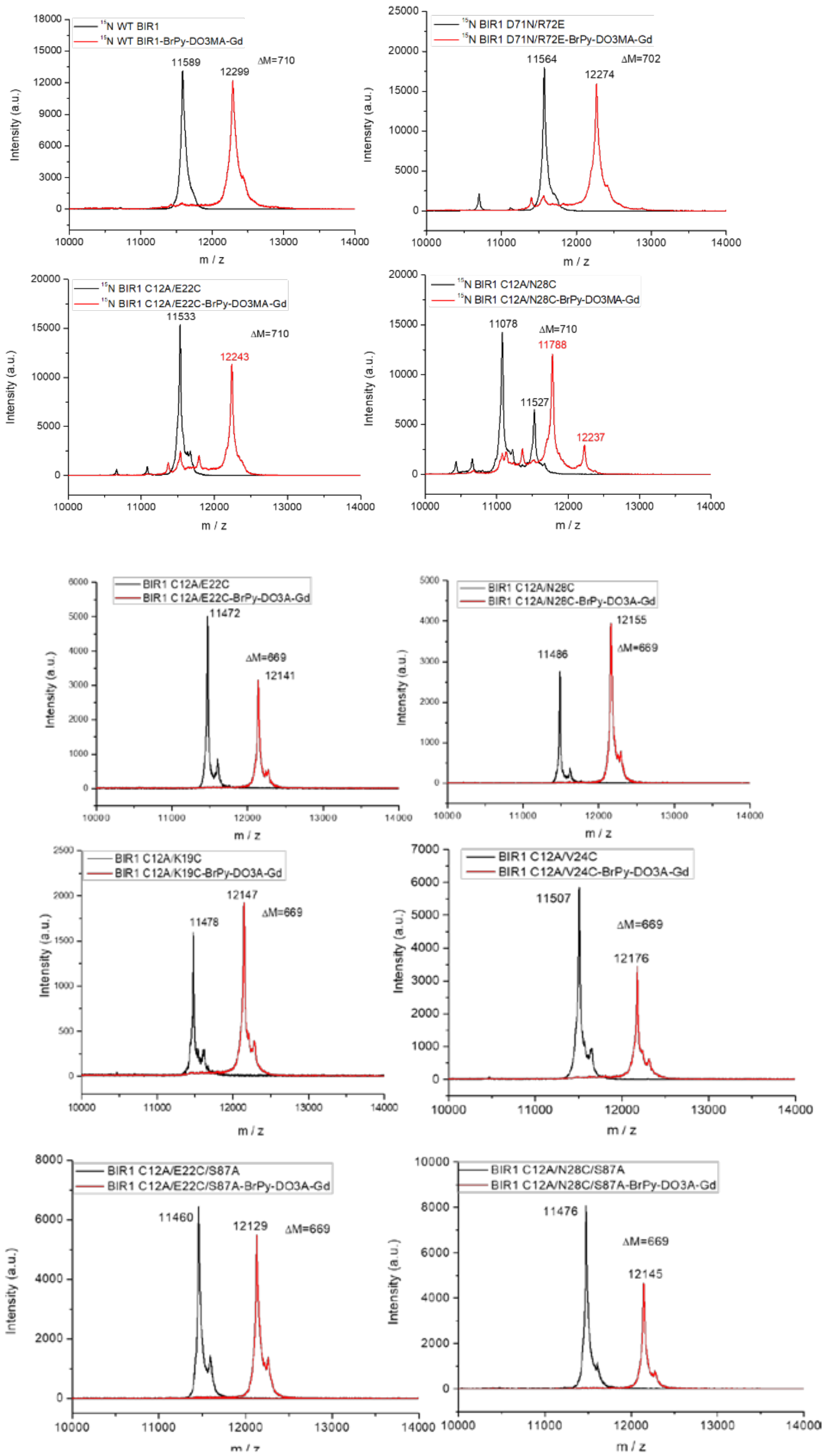


**Figure S1.** MALDI-TOF mass spectra of $^{15}$N labeled WT BIR1 and its mutant before (black) and after labeling with **GdII** or **GdI** complex (red), as noted on the Figure. The observed mass difference between non-labeled protein and labeled protein in a monomeric subunit is consistent with theoretical mass difference, 710 and 669, respectively, indicating labeling of only one site. It is noted that the peak corresponding to the molecular weight of $^{15}$N labeled BIR1 C12A/N28C, 11527, is not the most intense peak in the mass spectra. The and the integrity of the protein was confirmed by $^{15}$N-HSQC as shown in Figure S3.

### 3. NMR measurements

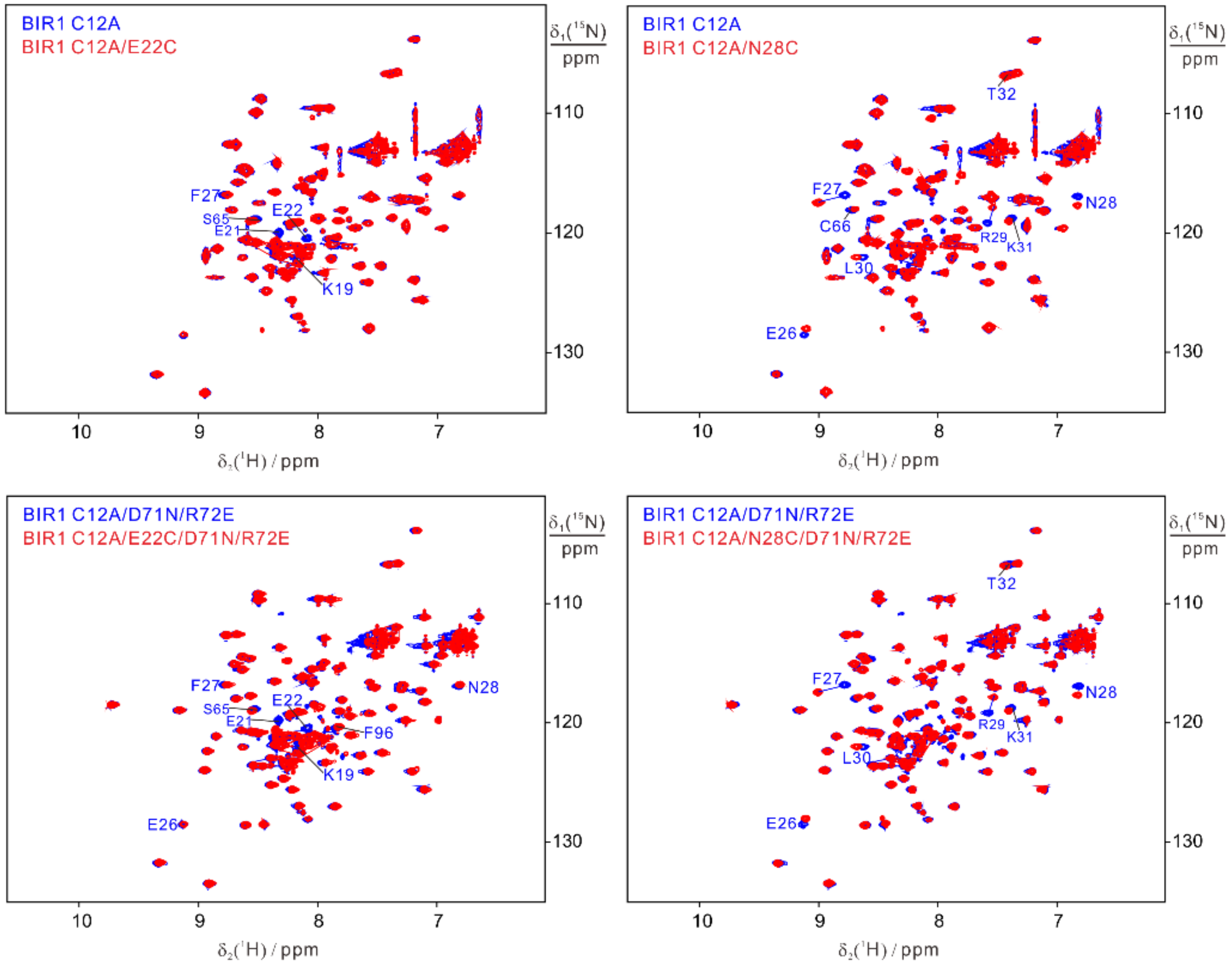


**Figure S2.** Superimposition of $^{15}$N-HSQC spectra recorded for 0.15 mM $^{15}$N labeled WT BIR1 (blue) and its cysteine mutant (red), indicating that the cysteine mutation did not change the overall chemical shift perturbations. All the NMR experiments were performed in 20 mM Bis-Tris, pH 6.5, and at 298K.

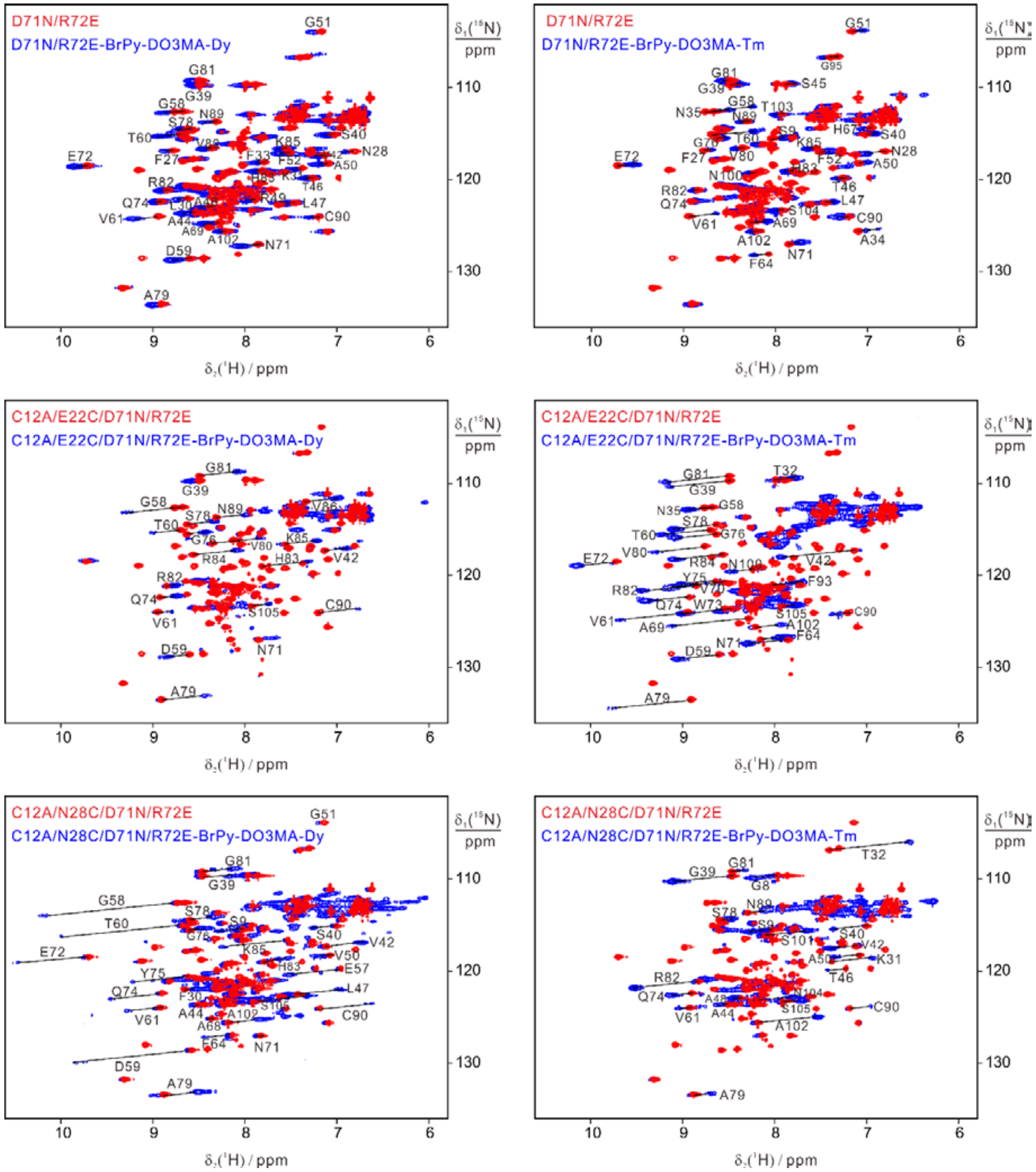


**Figure S3.** Superimposition of $^{15}$N-HSQC spectra recorded for 0.15 mM $^{15}$N labeled BIR1 D71N/R72E or its cysteine mutants (red) and the corresponding BrPy-DO3M(R)A-Ln (Ln = $Dy^{3+}$ or $Tm^{3+}$) conjugate complex (blue), for D71N/R72E (top panel), C12A/E22C/D71N/R72E (middle) and C12A/N28C/D71N/R72E (bottom). All the NMR experiments were performed in 20 mM Bis-Tris, pH 6.5, and at 298K.

The flexibility gradient within C12, E22 and N28 was determined from paramagnetic NMR measurements, which were carried out on the $^{15}$N-enriched monomeric D71N/R72E, C12A/E22C/D71N/R72E, and C12A/N28C/D71N/R72E BrPSPy-DO3MA-Ln (Ln = $Dy^{3+}$ and $Tm^{3+}$). This was justified because dimeric WT BIR1 and monomeric D71N/R72E have an almost identical 3D structural fold and it is more straightforward to assess the PCSs in a monomeric state. A single point mutation (E22C or N28C) did not change the overall protein structure, as indicated by mapping of the chemical shifts in the $^{15}$N-HSQC spectra (Figure S2). The D71N/R72E–

BrPy-DO3MA-Ln (Ln = $Dy^{3+}$ and $Tm^{3+}$) conjugates showed small PCSs in their $^{15}$N-HSQC spectra, whereas the C12A/E22C/D71N/R72E–BrPy-DO3MA-Ln and C12A/N28C/D71N/R72E–BrPy-DO3MA-Ln conjugates showed markedly larger PCSs (Figure S3).

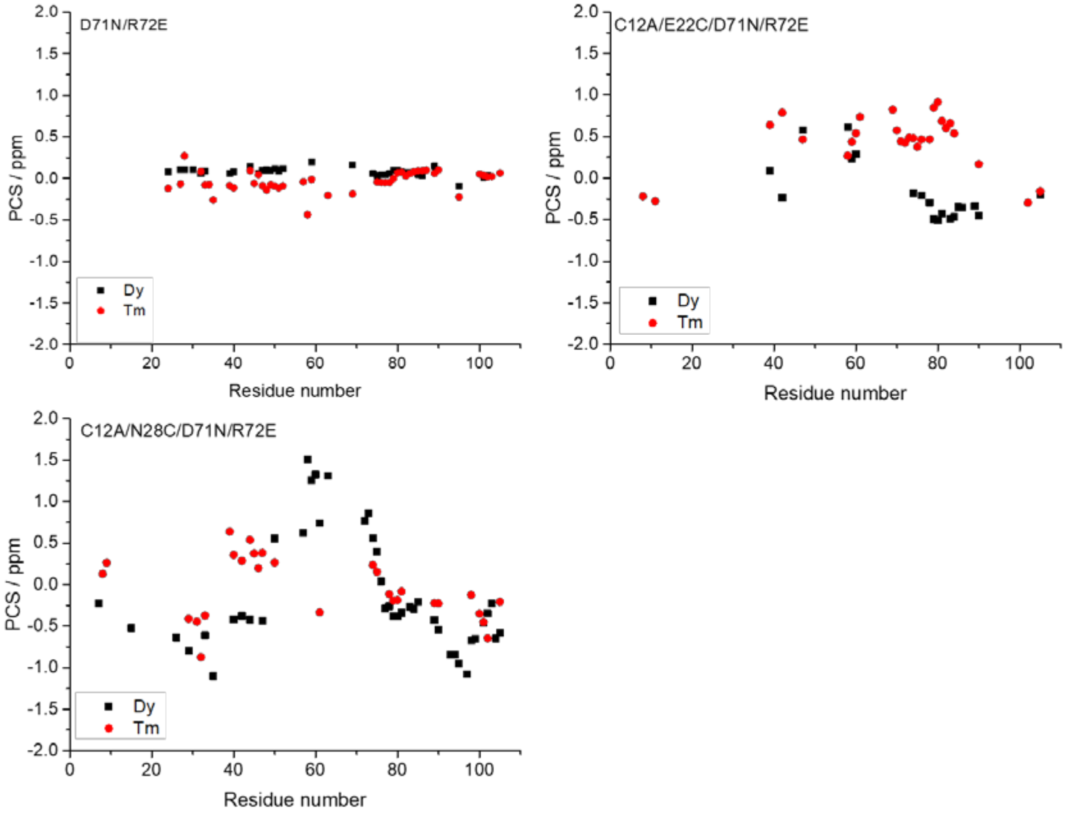


**Figure S4.** Plot of PCSs of backbone amide protons, determined for the monomeric BIR1-BrPy-DO3MA-Ln (Ln = $Tm^{3+}$ or $Dy^{3+}$) complex, as a function of amino acid sequence.

### 4. DEER data of monomeric BIR1

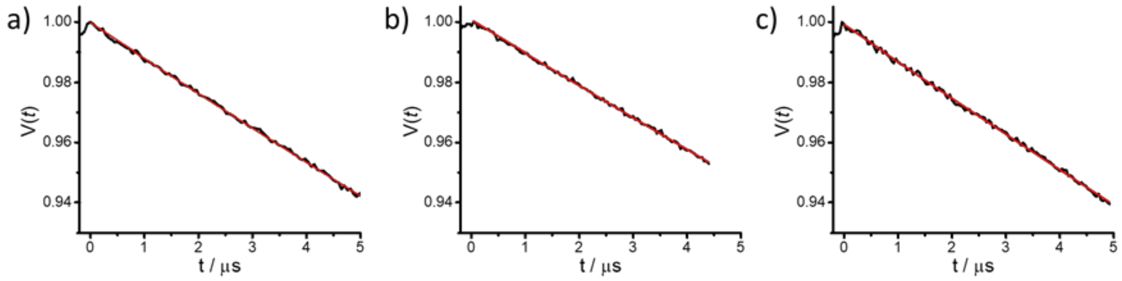


**Figure S5.** Primary DEER traces of 200 μM BIR1 conjugates in buffer solution. (a) D71N/R72E/C12-**GdII**, (b) D71N/R72E/C12A/N28C- **GdII** and (c) S87E/C12-**GdII**.

### 5. EPR and echo-decays

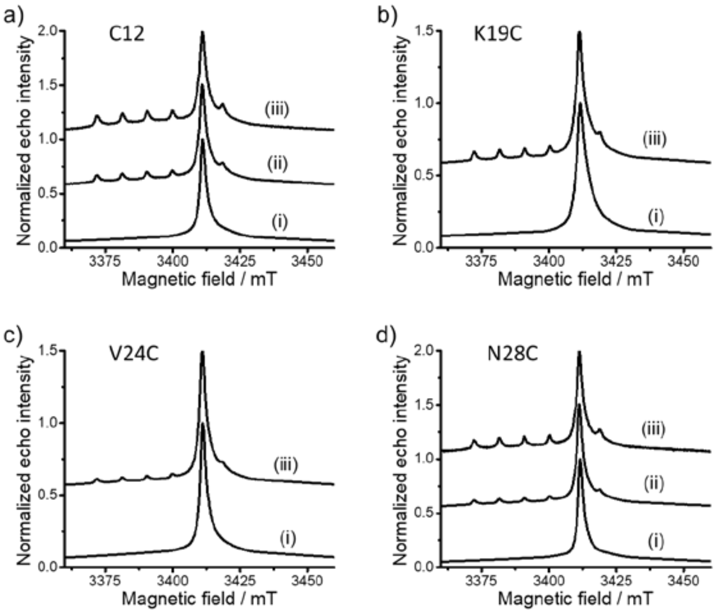


**Figure S6.** The central transition region of the W-band ED-EPR spectra of BIR1 constructs labeled with **GdI** different environments as noted on the Figure, where C12, K19C, V24C and N28C denote WT BIR1, C12A/K19C, C12A/V24C and C12A/N28C, respectively. (i) 200 μM of BIR1 conjugates in 20 mM Tris (pD 7.2). (ii) 30 μM of BIR1 conjugates in cell lysate. (iii) BIR1 conjugates in HeLa cells, frozen 5 hours after delivery by electroporation. Spectra were normalized to unity and are shifted by 0.5 to ease comparison.

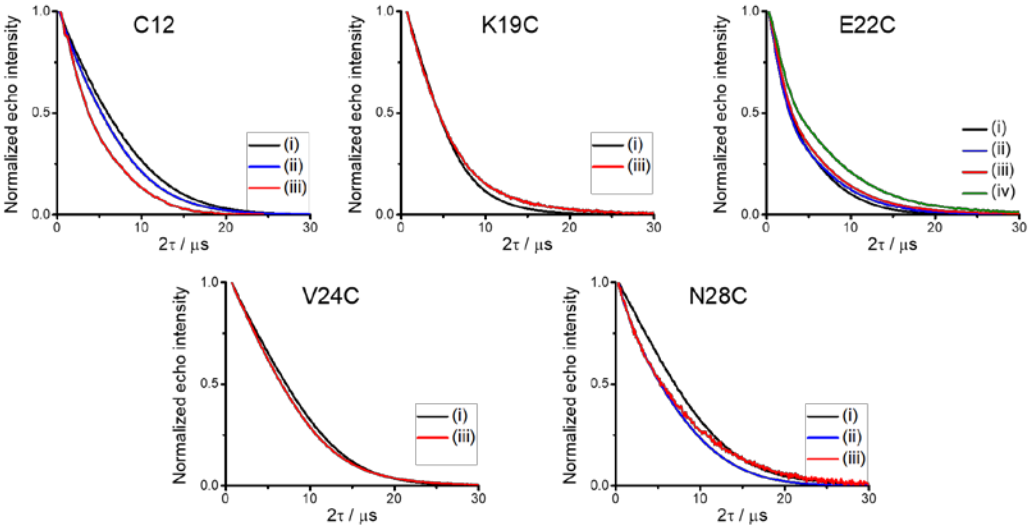


**Figure S7.** Two pulse-echo decay of BIR1 constructs labeled with **GdI** in different environment. (i) 200 μM of BIR1 conjugates in 20 mM Tris (pD 7.2). (ii) 30 μM of BIR1 conjugates in HeLa cell lysate. (iii) BIR1 conjugates in HeLa cells, frozen 5 hours after delivery by electroporation. (iv) Lysate of the electroporated cells.

### 6. Primary DEER data and analysis for BIR1 mutants

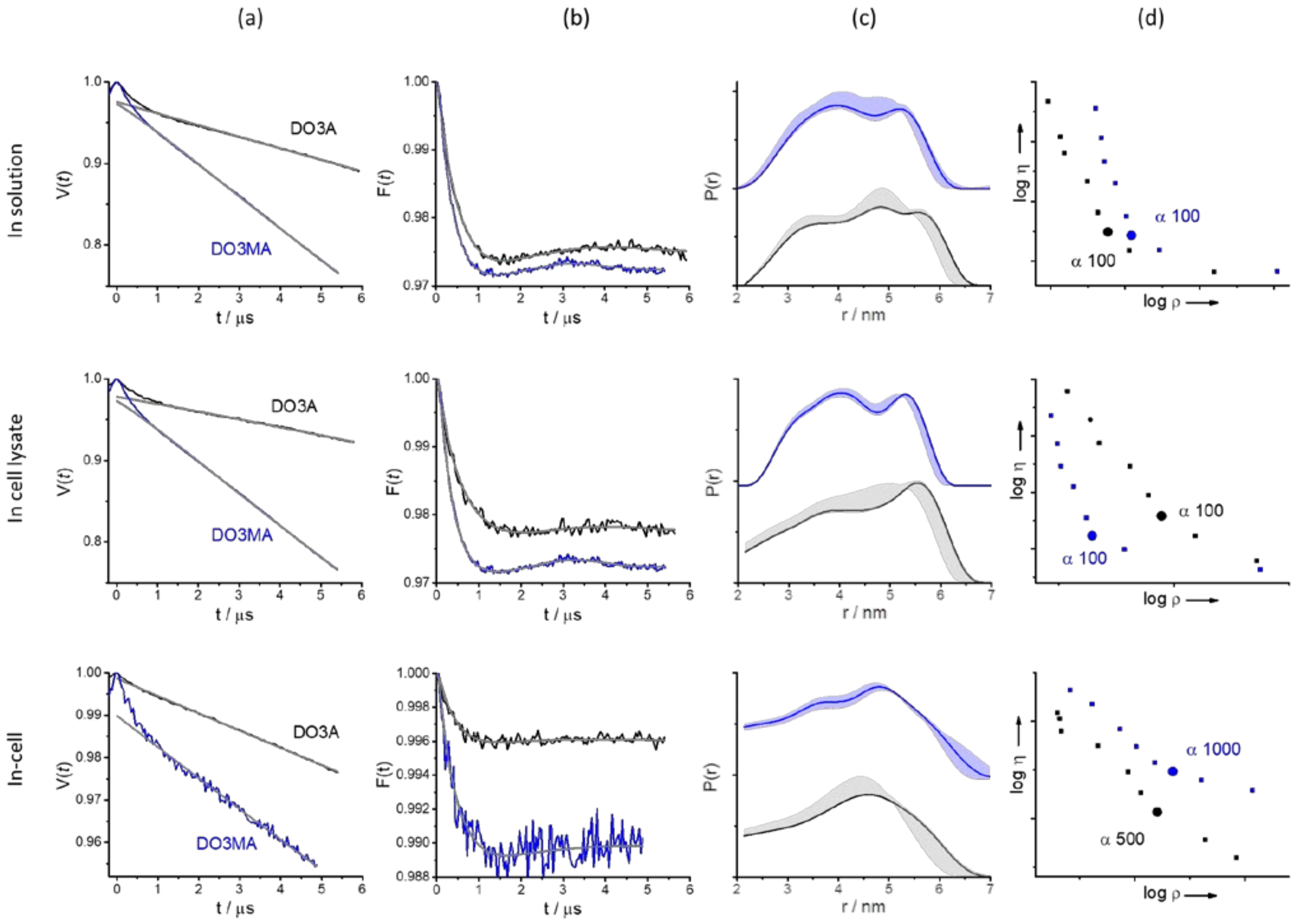


**Figure S8.** DEER results of BIR1 C12-**GdI** (marked as DO3A) and BIR1 C12 -**GdII** (marked as DO3MA) in different environments: in 20 mM Tris (pD 7.2), in cell lysate and in HeLa cells frozen after incubation for 5 hours at 37 ºC after electroporation delivery. (a) Primary DEER data with the background decay function in grey. (b) The DEER form factor after background removal and the fit (gray line) obtained with the distance distribution indicated in (c). (c) The distance distribution and the uncertainty obtained through the validation option in DeerAnalysis. (d) The corresponding L-curve produced with Tikhonov regularization, the chosen α-value is indicated on the figure as a larger symbol. The background slope and modulation depth in DEER traces of **GdII** labeled samples are generally bigger than those of the corresponding **GdI** labeled samples. This is due to the different labeling efficiencies of these two tags to BIR1 mutants, as mentioned in the experimental section.

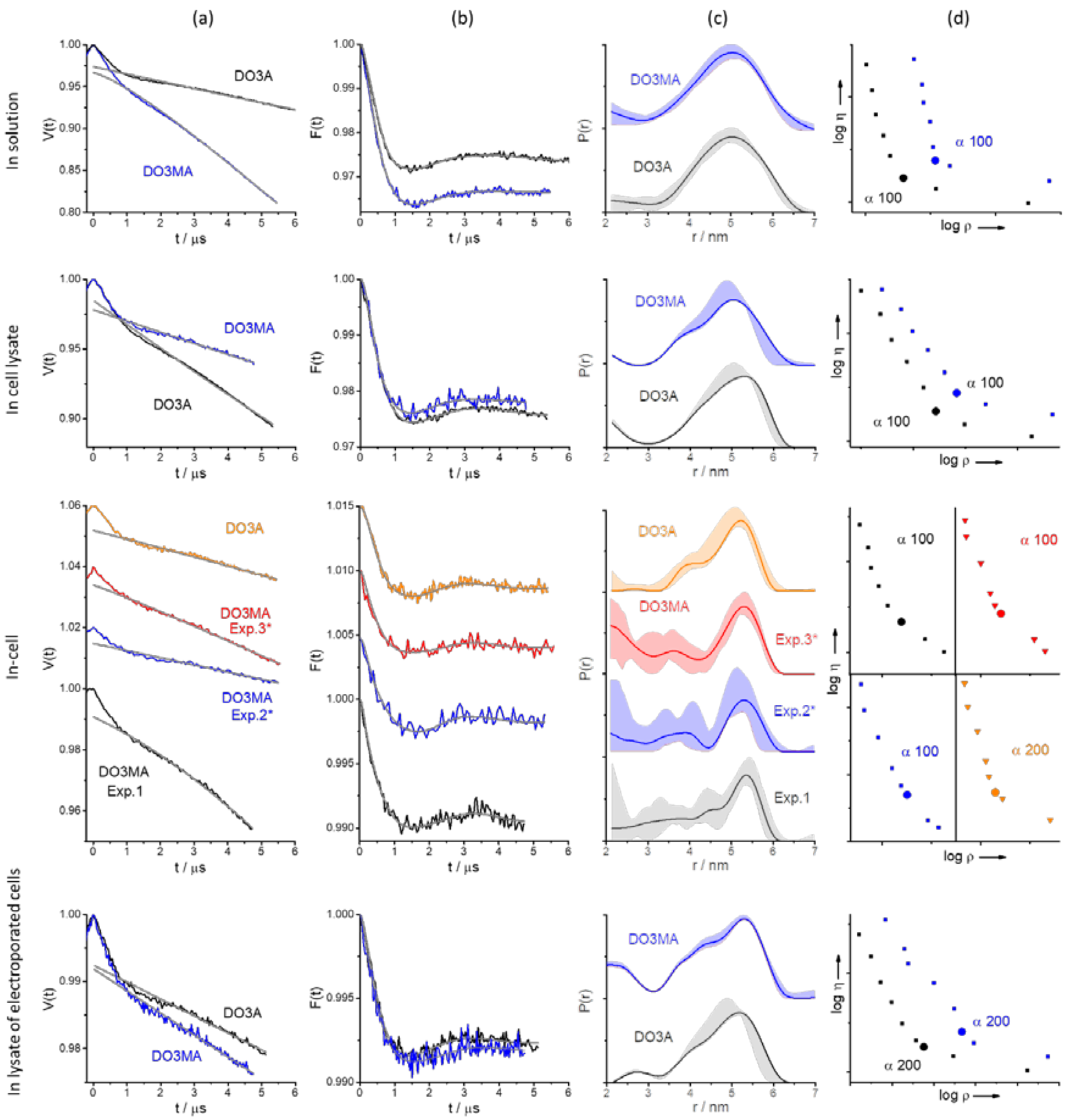


**Figure S9.** DEER results of BIR1 C12A/E22C- **GdI** (marked as DO3A) and C12A/E22C- **GdII (**marked as DO3MA) in different environments as noted on the figure. * marks DEER traces measured with rectangular pump pulse. The a, b, c, d notations are as in Figure S8.

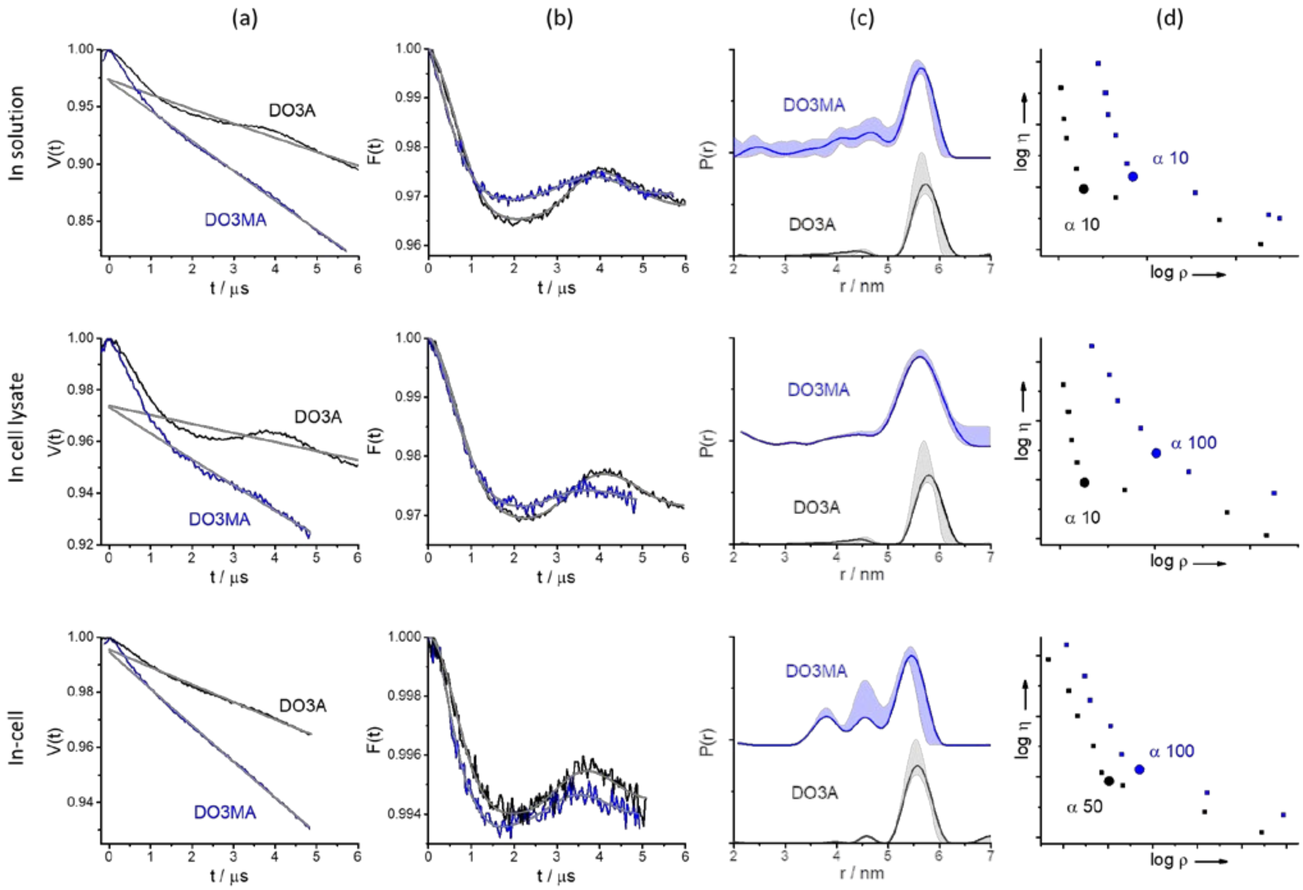


**Figure S10.** DEER results of BIR1 C12A/N28C-**GdI (**marked as DO3A, black**)** and C12A/N28C-**GdII (**marked as DO3MA, blue**)** in different environments as noted on the figure. The a, b, c, d notations are as in Figure S11.

### 7. Primary DEER data and analysis for E22C and N28C in the presence of co-solutes

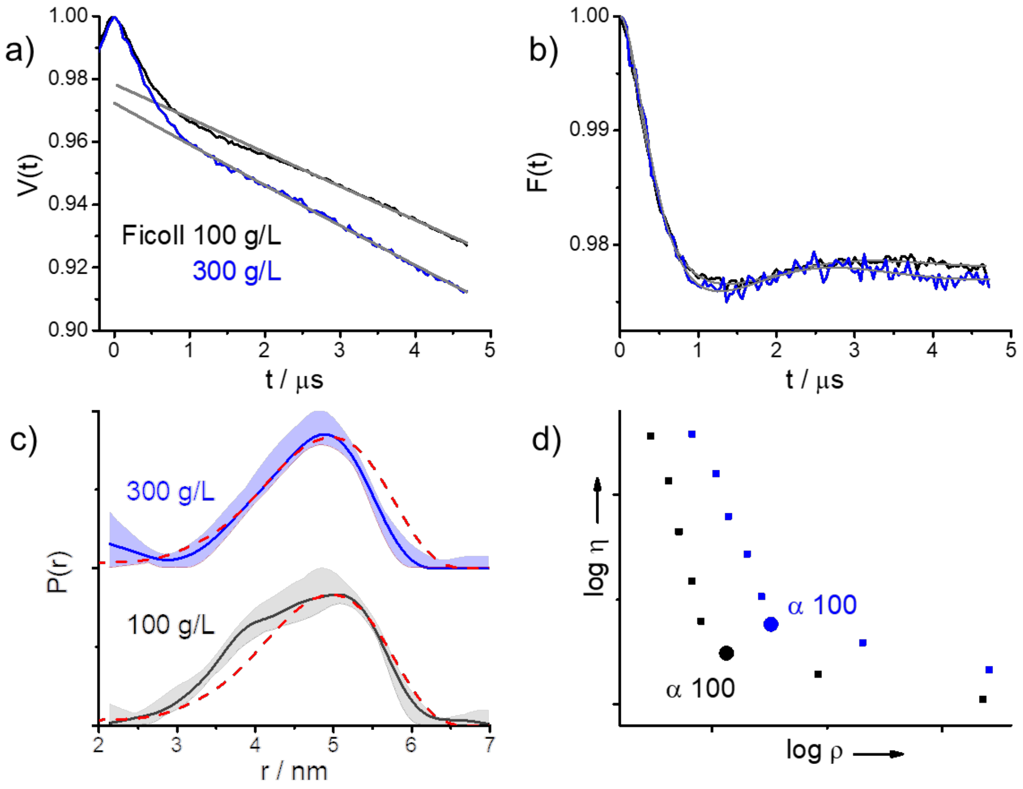


**Figure S11.** DEER results of 200 μM BIR1 C12A/E22C-**GdI**in the presence of Ficoll in 20 mM Tris (pD 7.2). (a) Primary DEER data with the background decay function in gray. (b) The DEER form factor after background removal and the fit (gray line) obtained with the distance distribution indicated in (c). (c) The distance distribution and the uncertainty obtained through the validation option in DeerAnalysis. The dashed red lines correspond to the distance distribution without Ficoll. (d) The corresponding L-curve produced with Tikhonov regularization. The chosen α-value is indicated on the figure.

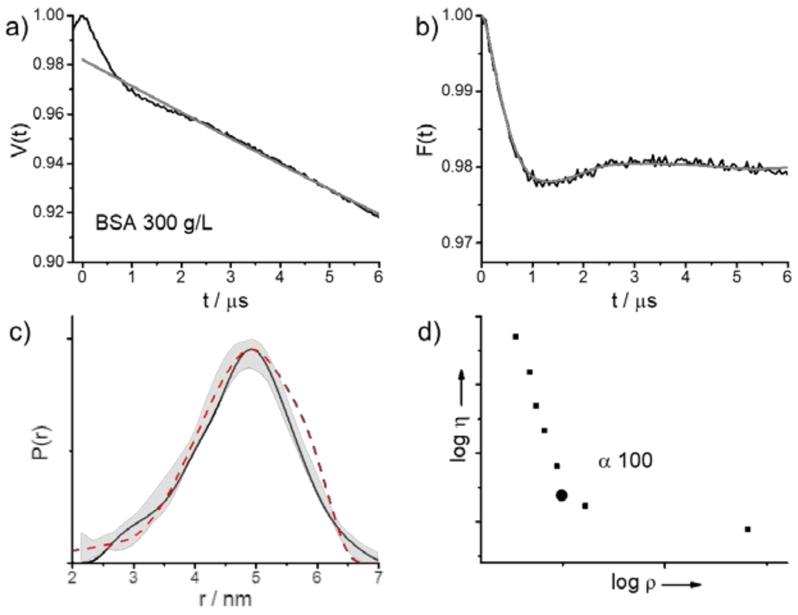


**Figure S12.** DEER results of BIR1 C12A/E22C-**GdI** in the presence of BSA in 20 mM Tris (pD 7.2). The red dashed line in (c) corresponds to the distance distribution without BSA. The a, b, c, d notations are as in Figure S8.

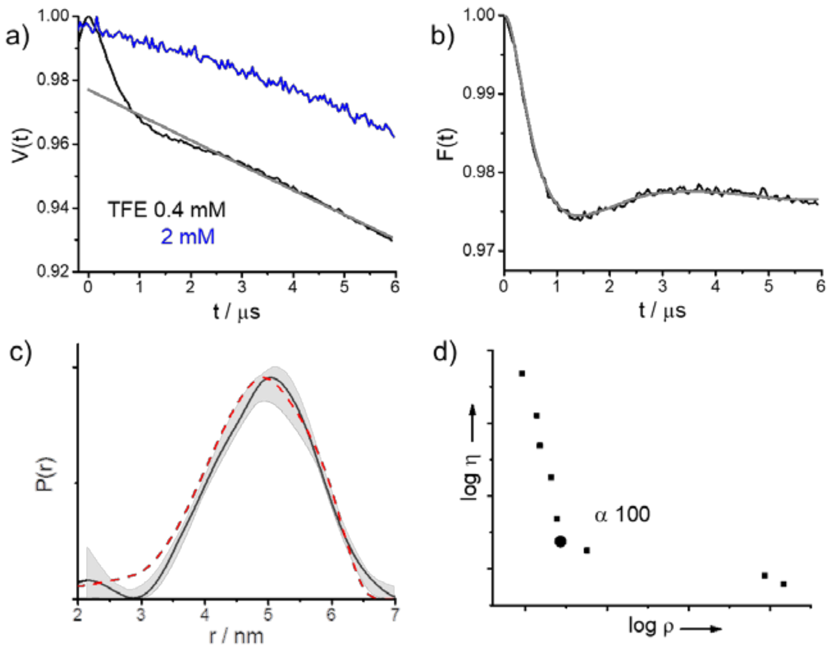


**Figure S13.** DEER results of BIR1 C12A/E22C-**GdI** in the presence of TFE (Trifluoroethanol) in 20 mM Tris (pD 7.2). The red dashed line in (c) corresponds to the distance distribution without TFE. The notation a, b, c, d notations are as in Figure S8

### 8. Primary DEER data and analysis for K19C and V24C

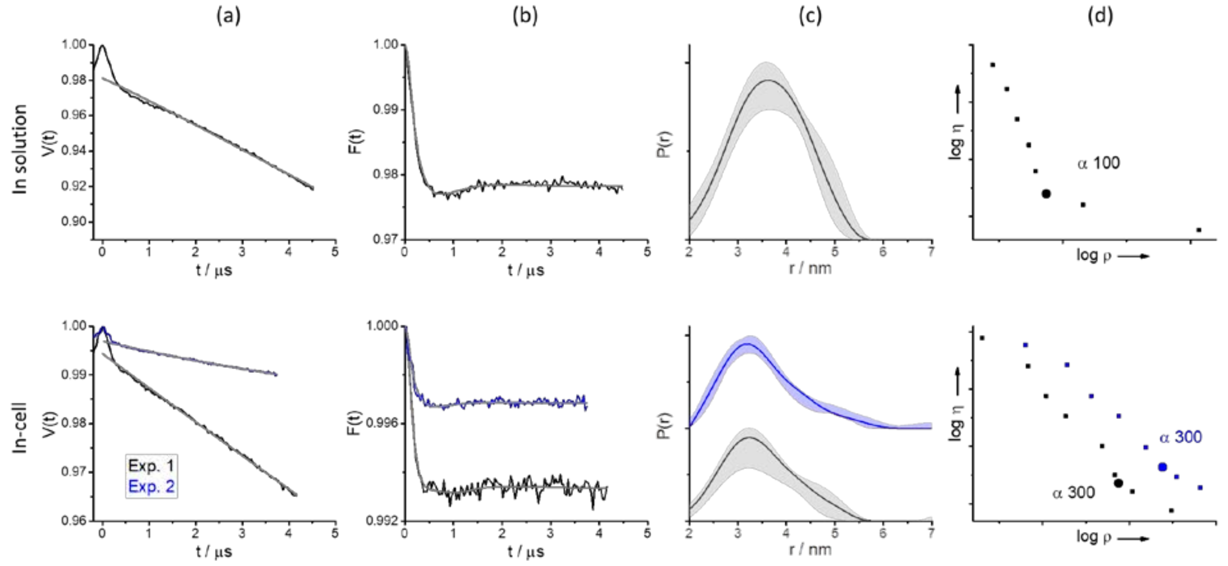


**Figure S14.** DEER results of BIR1 C12A/K19C-**GdI** in solution (top raw) and in cells (bottom raw).The a, b, c, d notations are as in Fig. S8.

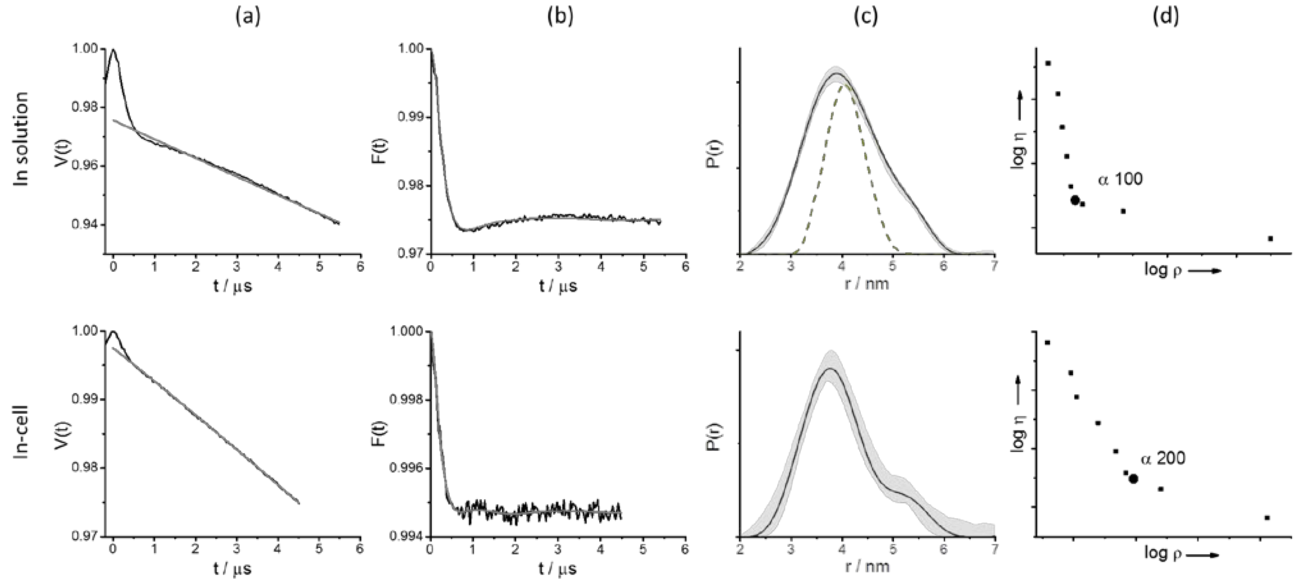


**Figure S15.** DEER results of BIR1 C12A/V24C-**GdI** in solution (top raw) and in cells (bottom raw). The a, b, c, d notations are as in Fig. S8. The dashed trace in c corresponds to the predicted distance distribution Distance distributions predicted by MtsslWizard.

### 9. Determination of dissociation constants from DEER data

The dissociation constants, $K_D$, of dimeric BIR1 conjugates were determined from the modulation depth, λ, of in-solution DEER form factors (all recorded under the same conditions) on a series of **Gd**I-labeled C12A/N28C and C12A/S87A/N28C conjugates at various concentrations in 20 mM Tris solution (pD 7.2). For the former we also included in the data the solution λ values of **GdI** conjugates of WT C12 and C12A/E22C. The DEER traces were analyzed by fitting of the background decay with three-dimensional homogenous distribution (exponential decay) taking into account the time window between 2200 and 5800 ns. In practice, because of the low λ, the background decay was linear.

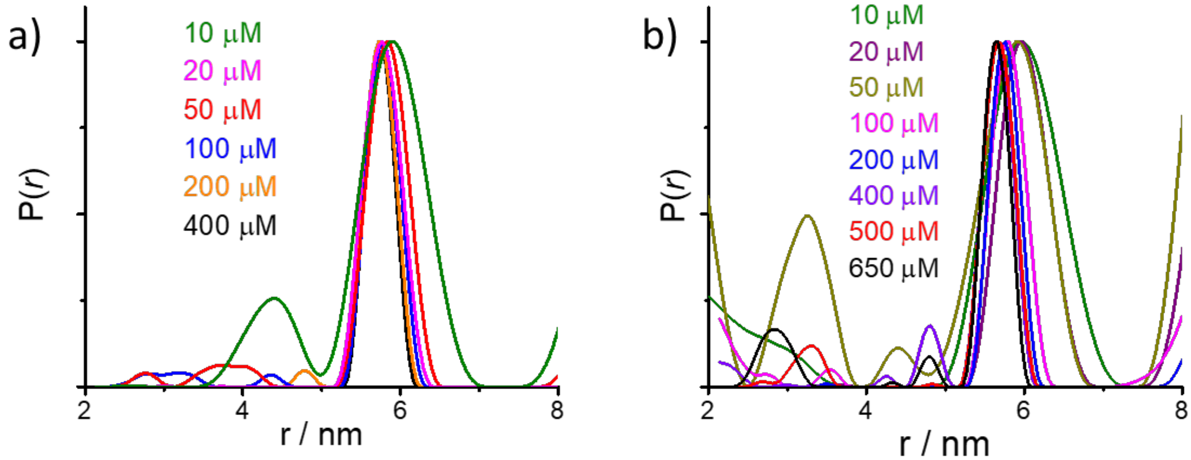


**Figure S16.** The distance distributions obtained from DEER results of BIR1 conjugates in at various concentrations (noted on the figure) in 20 mM Tris (pD 7.2), shown in Fig. 4 and Fig. 5 for (a) BIR1 C12A/N28C-**GdI**. (b) S87A/C12A/N28C-**GdI** , respectively.

### 10. Monomer-dimer equilibrium observed by NMR

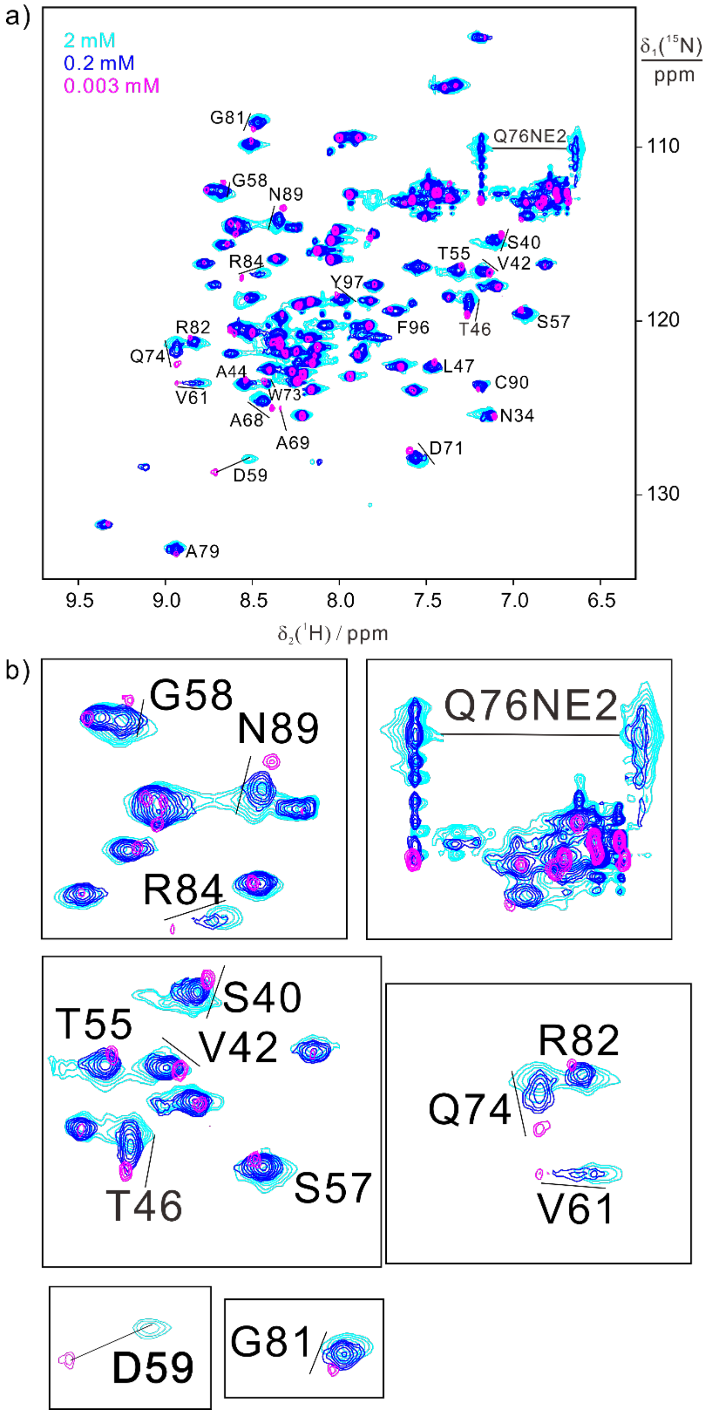


**Figure S17.** a) Superposition of $^{15}$N-HSQC spectra of $^{15}$N-labeled WT BIR1 at different concentrations, 2 mM (cyan), 0.2 mM (blue), and 0.003mM (magenta), in 20 mM Bis-Tris, pH 6.5, and at 298K. The residues experiencing chemical shift changes as a function of protein concentrations are labeled. b) Zoomed regiosn in a).

### 11. SEC results on WT BIR1, S87A and S87E mutants

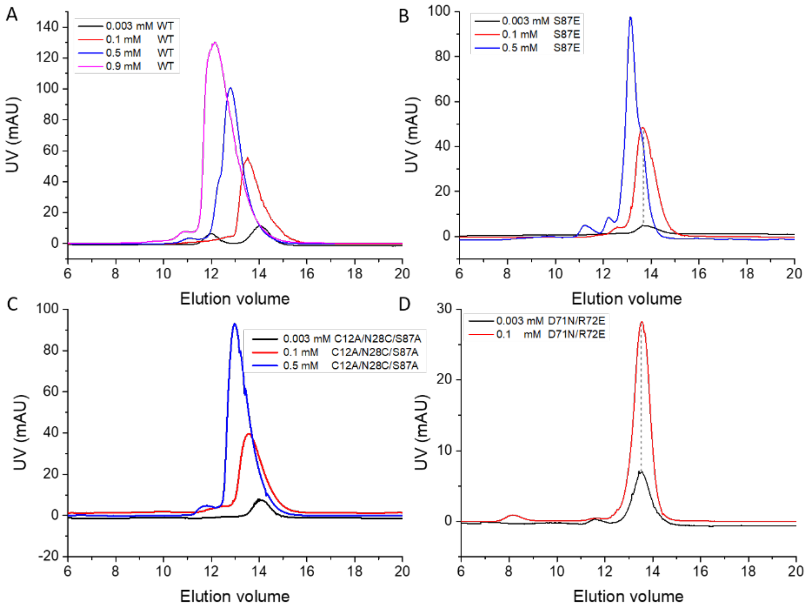


**Figure S18**. Gel filtration profiles of WT BIR1 and its mutants at various concentrations as noted on the figure, of which the absorption was monitored by UV lamp at 280 nm.

### 12. NMR relaxation data on WT BIR1, D71N/R72E, S87A and S87E mutants

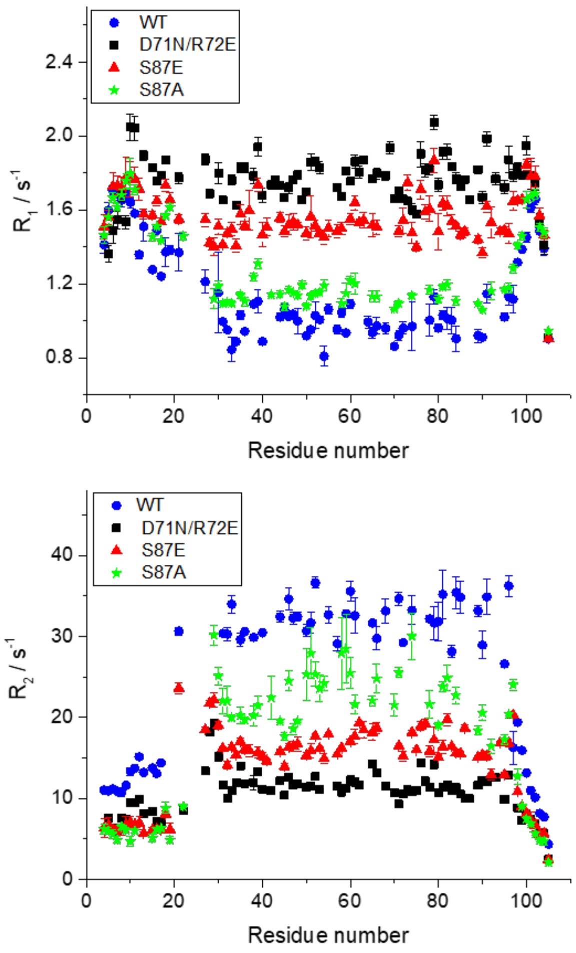


**Figure S19.** Comparison of backbone $^{15}$N-relaxation rates measured for 0.3 mM WT BIR1 (black), 0.4 mM D71N/R72E (red), 0.5 mM S87E (blue) and 0.5 mM S87A mutants at 298 K in 20 mM Bis-Tris buffer at pH 6.5 with a proton frequency of 600 MHz. The data of WT BIR1 and D71N/R72E were reported in the previous study.[1] It is evident that the transverse relaxation rates, $R_2$, decrease from WT BIR1, S87A, S87E to D71N/R72E, suggesting that the averaged molecular weight in solution decreases following WT BIR1, S87A, S87E and D71N/R72E.

### 13. Determination of in-cell concentrations

The background decay for a homogenous three dimensional distribution of spins is given by (11)

$$B(t) = V_0 e^{-kC\lambda'/t} \qquad \text{(S1)}$$

where $V_0$ is the echo intensity at $t$=0, $C$ is the spin concentration in the sample , $\lambda$' is the modulation depth , namely fraction of spins inverted by the pump pulse, and $k$ is a constant

$$k = \frac{2\pi}{9\sqrt{3}}\frac{g^2\mu_B^2\mu_0}{\hbar} = 0.997 \quad \text{(S2)}$$

in units of $\mu s^{-1}$ $mM^{-1}$. A plot of *lnB(t)* vs *t* generates a straight line with a slope $b = -kC\lambda'$ and a plot of *b* vs *C* yields yet another linear relationship with a slope $d= -k\lambda'$ from which $\lambda'$ can be determined (12). Unlike the intra-dimer modulation depth, $\lambda$, used in the main text, which depends on the fraction of dimers in the system and is therefore concentration dependent, *b* depends on the total spin concentration in sample and $\lambda'$ depends on the pump pulse parameters and the spectral lineshape and is concentration independent. In the case of $Gd^{3+}$ *b* is small and *B(t)* is linear. When the labeling efficiency, *f*<0, the protein concentration, $C_P$ and the spin concentration are related by $C=fC_p$.

We have generated calibration curves of *b* vs $C_P$ (Figs. 4c and 5c) from which the local in-cell concentration can be determined, once the in-cell measurements are carried out under the same conditions as the solution ones such that $\lambda'$ remains constant.

The in-cell $C_p$ obtained using the calibration curves in Figs. 4c and 5c are overestimated because of the presence of endogenous $Mn^{2+}$, which contributes to the background decay. In this case one has to take into account the dipolar interaction of $Gd^{3+}$) with both the $Gd^{3+}$ and Mn(II) spins in the sample. Because we set the observe pulse to the maximum of the $Gd^{3+}$signal, which is considerably higher than that of the $Mn^{2+}$at this field position (see Fig. 2) we can neglect the $Mn^{2+}$ contributions to the observed signal and consider only its contribution to the flipped spin by the pump pulse. In this case we write

$$b = -k(\lambda'_{Mn}C_{Mn} + \lambda'_{Gd}C_{Gd}) \text{ (S3)}$$

Where $\lambda'_{Mn}$ and $\lambda'_{Gd}$ correspond to the probability of the pump pulse to invert a $Gd^{3+}$ and a $Mn^{2+}$spin respectively and $C_{Mn}$ and $C_{Gd}$ are their corresponding concentrations in the sample. From eq. S3 we obtain:

$$b = -k\lambda'_{Gd}C_{Gd}\left(1 + \frac{\lambda'_{Mn}C_{Mn}}{\lambda'_{Gd}C_{Gd}}\right) = -k\lambda'_{Gd}C_{Gd}(1 + x) \quad \text{(S4)}$$

And $x = \frac{\lambda_{Mn}C_{Mn}}{\lambda_{Gd}C_{Gd}}$ is the correction factor. Accordingly, *b* is overestimate by a factor of *(1+ x)*, and the same goes for C derived from the measured *b* and the corresponding protein concentration $C_p=C/f$ (red points in Fig. 4c and 5d) Therefore, we have to determine *x* in order to obtain the correct in-cell protein contribution.

The in-cell spectrum is a superposition of the spectrum of the endogenous $Mn^{2+}$ (Fig. S21a) and that of the labeled protein with BrPy-DO3A-Gd given in Fig. S21b. To determine $\frac{C_{Mn}}{C_{Gd}}$ we simulated the experimental ED-EPR spectra as a superposition of these two spectra as with their areas normalized to 1 and multiplying the $Gd^{3+}$ spectrum by 4/3, which to a good approximation represents its higher transition probabilities. The relative weights of the two spectra in the final fitted spectrum is $\frac{C_{Mn}}{C_{Gd}}$ and the values obtained are listed in Table S1. Next we evaluated $\frac{\lambda'_{Mn}}{\lambda'_{Gd}}$ by calculating the area covered by the chirp pump pulses for each of the spectra relative to the area of the full spectra. While the chirp pulses have a width of 300 MHz, the affected area is limited to no more than 100 MHz because of the width of the cavity. This generated for $\lambda_{Gd}$=7.6% and $\lambda_{Mn}$=3.7% and $\frac{\lambda'_{Mn}}{\lambda'_{Gd}}$ =0.48. In Table S1 we list the corrected in cell concentrations for the various constructs along with the predicted $\lambda$ for the i*n-vitro* $K_D$.

We note that under our measurements conditions, with the observe pulse set to the maximum of the $Gd^{3+}$ signal the $Mn^{2+}$ contribution to the background decay is significant but to λ it is small. The situation is opposite when the pump pulse is set to the maximum of the $Gd^{3+}$ signal, and the observe outside this transition.

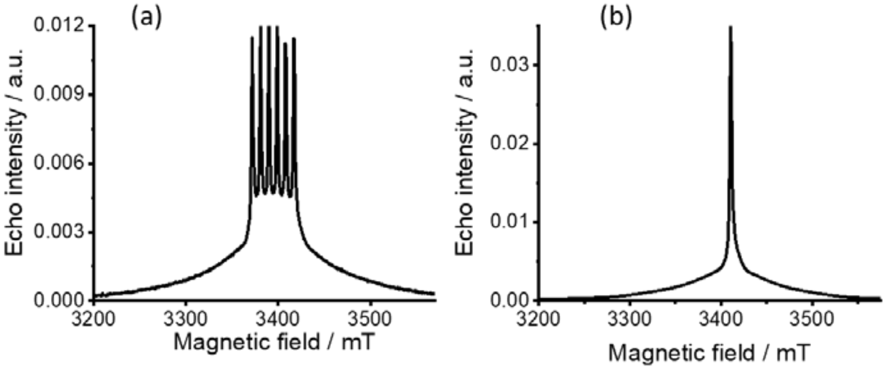


**Figure S20.** W-band ED-EPR (10 K) spectra of Hela cells and (b) BIR1 C12A/N28C-**GdI** in solution.

**Table S1.** Summary of the DEER modulation depth obtained for BIR1 constructs in vitro and in cell and the local protein concentration of in-cell samples.

| **GdI** labeled constructs | $\lambda$, in solution[a] | $\lambda$, in cell | in cell $C_p$ µM | $\frac{C_{Mn}}{C_{Gd}}$ | Corrected in cell $C_p$, µM | Predicted in cell $\lambda$ |
|---|---|---|---|---|---|---|
| C12 | 2.50±0.2 | 0.39±0.06 | 78.4±8 | 0.6±0.03 | 52.7±10 | 1.38±0.51 |
| C12A/K19C | 2.52±0.2 | 0.32 ±0.04 | 39.1±4 | 0.2±0.03 | 26.0±7.5 | 1.27±0.62 |
| | | 0.79 ±0.13 | 122.6±12 | 0.4±0.03 | 94.2±22.5 | 1.57±0.58 |
| C12A/E22C | 2.92±0.3 | 0.61 ±0.07 | 66.8±7 | 0.45±0.03 | 50.0±9.3 | 1.4±0.54 |
| C12A/V24C | 2.65±0.3 | 0.52± 0.6 | 96.5±10 | 0.2±0.03 | 78.6±20.5 | 1.31±0.58 |
| C12A/N28C | 2.84±0.3 | 0.50 ±0.06 | 115.1±15 | 0.4±0.03 | 84.5±16.4 | 1.30±0.54 |

[a]The protein concentration was 200 µM for all *in vitro* experiments except when indicated specifically.

### 14. DEER results on S87A/C12A/E22C-GdI and S87A/C12A/N28C-GdI

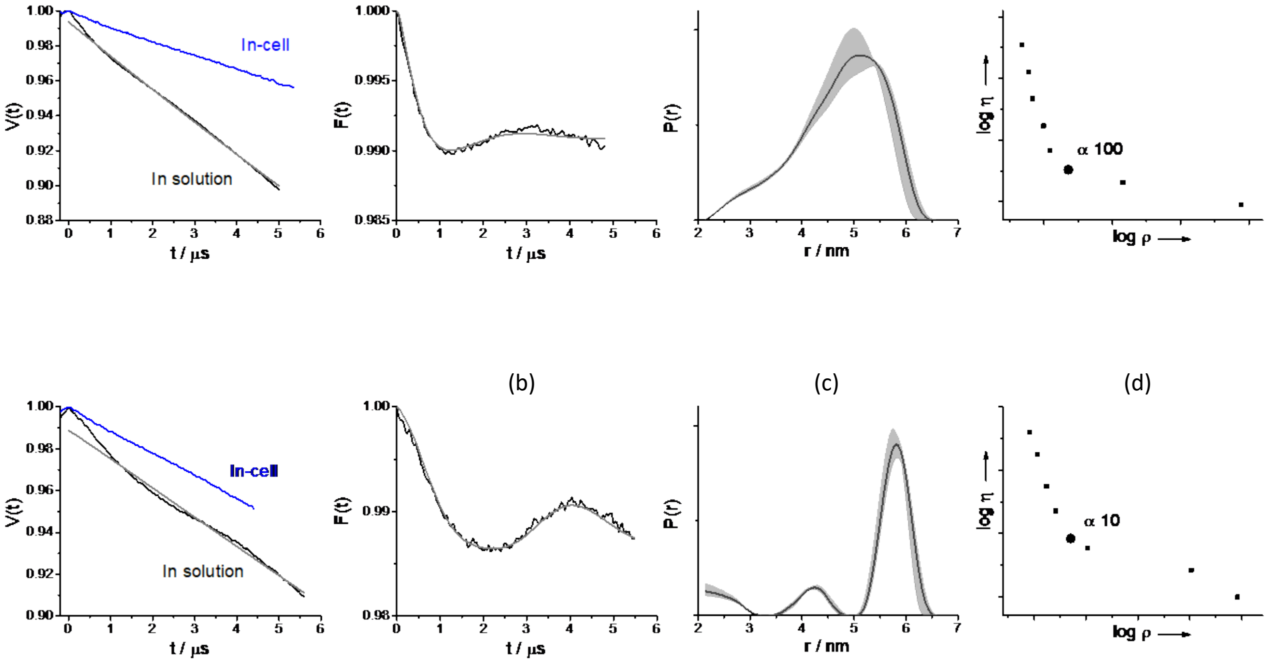


**Figure S20.** DEER results of BIR1 S87A/C12A/E22C-**GdI** (top raw) and BIR1 S87A/C12A/E22C-**GdI** (bottom raw) in different environments, as noted on the figure. The a, b, c, d notations are as in Fig. S8.

### 15. Effect of co-solutes on modulation depth

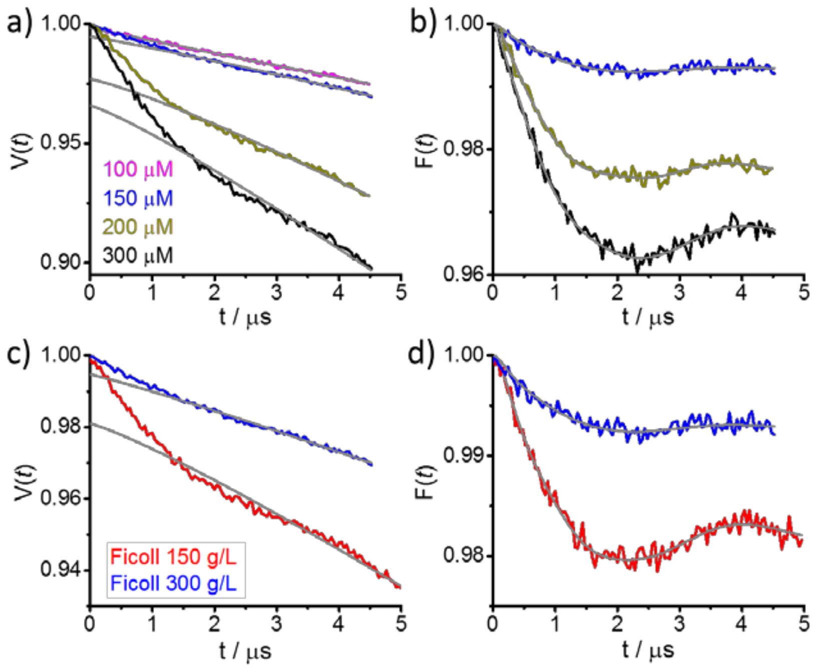


**Figure S21** . (a) Primary DEER traces with the background correction function DEER results for C12A/N28C–**GdI** in the presence of 300 g/L Ficoll in 20 mM Tris (pD 7.2) at various protein concentrations. (b) The corresponding form factors with the fit obtained from DeerAnalysis . (c) Primary DEER traces with the background correction function DEER results of 150 μM C12A/N28C–**GdI** in the presence of different amounts of Ficoll in 20 mM Tris (pD 7.2).

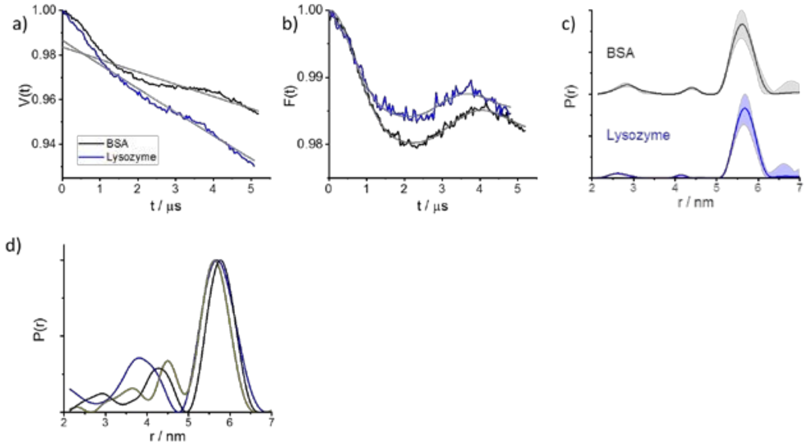


**Figure S22.** (a,b,c) DEER results of 100 μM C12A/N28C-**GdI** in the presence of BSA (300g/L) and lysozyme (20 g/L) in 20 mM Tris (pD 7.2). (d) Distance distributions of C12A/N28C- **GdI** in different concentrations in the presence of 300 g/L Ficoll in 20 mM Tris (pD 7.2). The DEER data are shown in Figure. S21a,b with the same color codes. The notation a, b, c, are as in Figure S8.

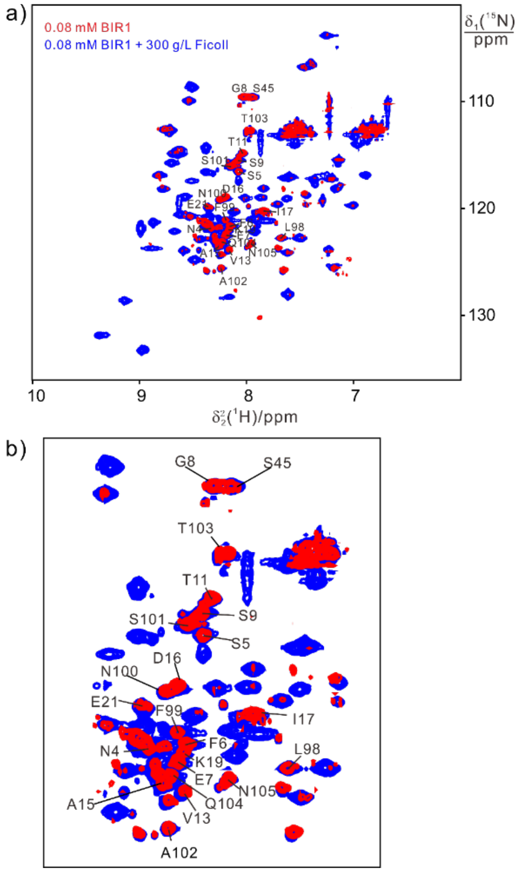


**Figure of S23**. a) Superimposition of $^{15}$N-HSQC spectra of 0.08 mM $^{15}$N-labeled WT BIR1 in the absence (blue) and presence of 300 g/L Ficoll (red). The spectra were recorded in 20 mM Bis-Tris, pH 6.5, and at 298K. b) Zoomed regions in a).

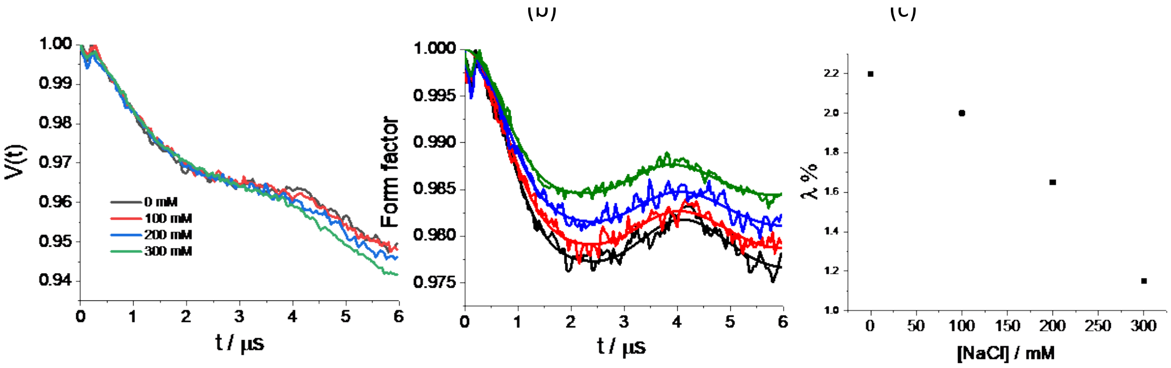


lys

**Figure 24.** DEER results for 100 µM C12A/N28C– GdI in the presence of salt. (a) Primary DEER traces, (b) DEER form factors after background removal using DeerAnalysis and (c) the dependence of λ on the NaCl concentration.

### 16. Electrostatic potential surfaces

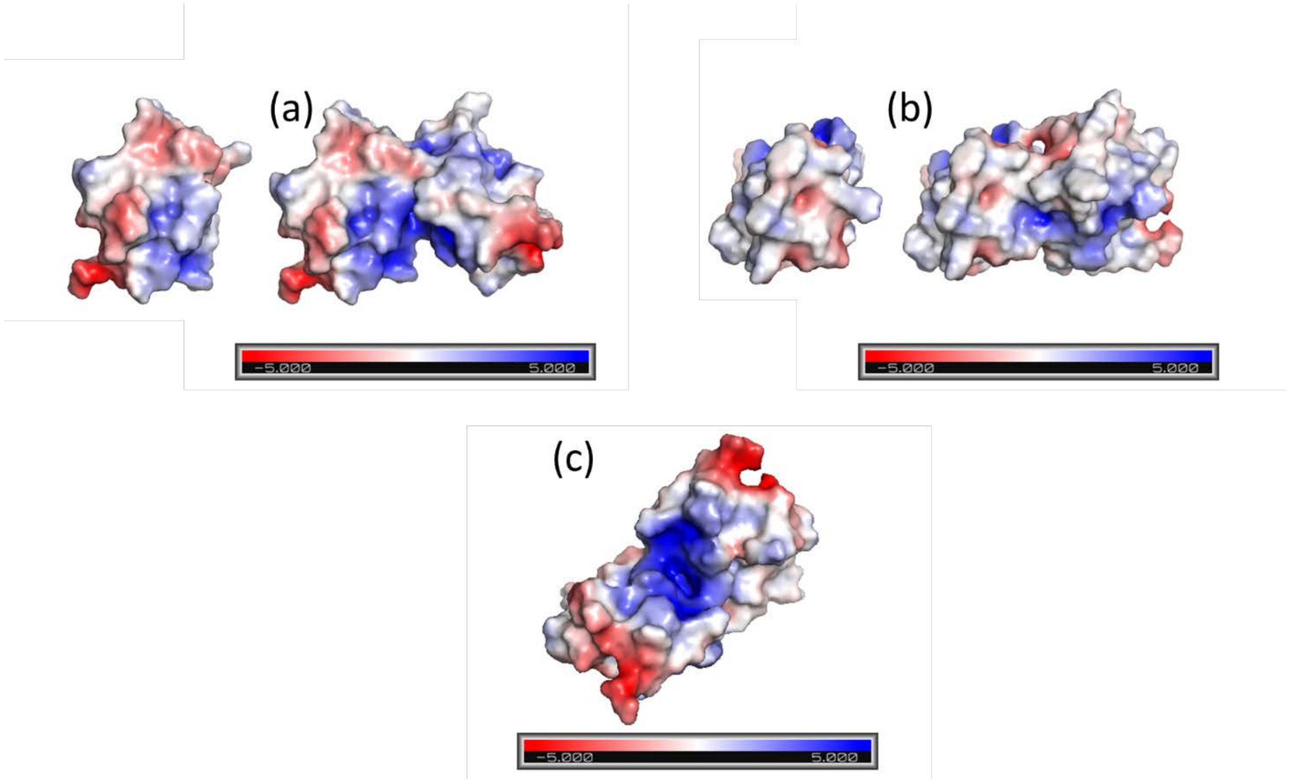


**Figure S25**. Electrostatic surface potential of the BIR1 monomer and dimer (a) and (b) at different orientations. In (c) the chosen orientation highlights the positive grove at the interface. The surfaces were obtained using the PyMOL plugin APBS electrostatics from the 2QRA crystal structure. The charge bar goes from -5.0 (red) to 5.0 (blue)

References

1. M. M. Hou *et al.*, Solution structure and interaction with copper in vitro and in living cells of the first BIR domain of XIAP. *Sci. Rep.* **7**, 16630 (2017).
2. Y. Yang *et al.*, A reactive, rigid Gd(III) labeling tag for in-cell EPR distance measurements in proteins. *Angew. Chem. Int. Ed. Engl.* **56**, 2914-2918 (2017).
3. Y. Yang *et al.*, High sensitivity in-cell EPR distance measurements on proteins using an optimized Gd(III) spin label. *J. Phys. Chem. Lett.* **9**, 6119-6123 (2018).
4. F. X. Theillet *et al.*, Structural disorder of monomeric alpha-synuclein persists in mammalian cells. *Nature* **530**, 45-50 (2016).
5. F. X. Theillet *et al.*, Site-specific NMR mapping and time-resolved monitoring of serine and threonine phosphorylation in reconstituted kinase reactions and mammalian cell extracts. *Nat. Protoc.* **8**, 1416-1432 (2013).
6. D. Goldfarb *et al.*, HYSCORE and DEER with an upgraded 95GHz pulse EPR spectrometer. *J. Magn. Reson.* **194**, 8-15 (2008).
7. F. Mentink-Vigier *et al.*, Increasing sensitivity of pulse EPR experiments using echo train detection schemes. *J. Magn. Reson.* **236**, 117-125 (2013).

8. Bahrenberg T., Yang Y., Goldfarb D., F. A., rDEER: A modified DEER sequence for distance measurements using shaped pulses. *Magnetochemistry* **5**, 20 (2019).M. Pannier, S. Veit, A. Godt, G. Jeschke, H. W. Spiess, Dead-time free measurement of dipole-dipole interactions between electron spins. *J. Magn. Reson.* **142**, 331-340 (2000).
9. G. Jeschke *et al.*, DeerAnalysis2006—a comprehensive software package for analyzing pulsed ELDOR data. *Appl. Magn. Reson.* **30**, 473-498 (2006).
10. A. M. Raitsimring, K. M. Salikhov, Electron-spin echo method as used to analyze the spatial distribution of paramagnetic centers. *Bull. Magn. Reson. 7*, 184-217 (1985).
11. S. Ruthstein, A. Potapov, A. M. Raitsimring, D. Goldfarb, Double electron electron resonance as a method for characterization of micelles *J. Phys. Chem. B 109*, 22843-51 (2005).
